# MERTK on mononuclear phagocytes regulates T cell antigen recognition at autoimmune and tumor sites

**DOI:** 10.1101/687061

**Authors:** Robin S. Lindsay, Jennifer C. Whitesell, Kristen E. Dew, Erika Rodriguez, Adam M. Sandor, Dayna Tracy, Seth F. Yannacone, Brittany N. Basta, Jordan Jacobelli, Rachel S. Friedman

## Abstract

Understanding mechanisms of immune regulation is key to developing immunotherapies for autoimmunity and cancer. We examined the role of mononuclear phagocytes during peripheral T cell regulation in type 1 diabetes and melanoma. MERTK expression and activity in mononuclear phagocytes in the pancreatic islets, promoted islet T cell regulation, resulting in reduced sensitivity of T cell scanning for cognate antigen in pre-diabetic islets. MERTK-dependent regulation led to reduced T cell activation and effector function at the disease site in the islets and prevented rapid progression of type 1 diabetes. In human islets, MERTK-expressing cells were increased in remaining insulin-containing islets of type 1 diabetic patients, suggesting that MERTK protects the islets from autoimmune destruction. MERTK also regulated T cell arrest in melanoma tumors. These data indicate that MERTK signaling in mononuclear phagocytes drives T cell regulation at inflammatory disease sites in peripheral tissues through a mechanism that reduces the sensitivity of scanning for antigen leading to reduced responsiveness to antigen.

## Introduction

Mechanisms of immune regulation function at all stages of the immune response to maintain and restore self-tolerance. These checkpoints provide layers of protection to control a normal immune response following pathogen clearance and to protect normal tissues from autoimmune destruction. These mechanisms of immune regulation also have the side effect of protecting many tumors from the immune response. The importance of understanding the multiple layers of immune regulation is highlighted by the fact that only 18%-36% of cancer patients respond to the targeting of a single checkpoint molecule (Hodi et al., 2010; Topalian et al., 2012) and response rates are improved by targeting multiple checkpoint molecules (Larkin et al., 2015). Thus, altering immune regulation for therapeutic purposes is likely to involve multiple pathways in most cancer and autoimmune patients.

Many layers of immune tolerance must be broken before autoimmune pathology occurs. Type 1 diabetes (T1D) is largely caused by the T cell mediated autoimmune destruction of the pancreatic islets. We have shown that the T cell response in the islets is temporally regulated. Antigenic restimulation of T cells occurs in early stages of islet infiltration leading to pathogenic T cell effector function, followed by suppression of T cell pathogenesis as islet infiltration advances (Friedman et al., 2014; Lindsay et al., 2015). This T cell regulation in the islets eventually fails in mice that present with overt T1D. The mechanisms that drive this temporally controlled regulation of T cell pathogenesis at the disease site in the islets are still mostly unknown. This period of T cell regulation is important, because it could represent a novel tolerance checkpoint in the islets during T1D progression. Establishing and maintaining tolerance in the islets following infiltration is a critical step towards halting disease progression after the autoimmune response has been initiated. Understanding the mechanism by which T cells are regulated in the islets during advanced infiltration must be accomplished before we can target this pathway to therapeutically protect tissues from ongoing autoimmune attack.

This period of T cell regulation in advanced infiltrated islets is associated with a loss of antigen-mediated T cell arrest and transient rather than sustained interactions with antigen presenting cells (APCs) (Lindsay et al., 2015; Friedman et al., 2014). T cell arrest and sustained interactions with APCs are generally associated with antigen recognition and stimulation, while T cell regulation or tolerance are associated with a lack of T cell arrest and with transient APC interactions (Jacobelli et al., 2013; Hugues et al., 2004; Katzman et al., 2010; Shakhar et al., 2005). For example, T cell tolerance through prevention of antigen-mediated T cell arrest has been shown to be mediated by factors such as regulatory T cells (Tadokoro et al., 2006; Tang et al., 2006) and co-inhibitory pathways including the PD-1 pathway (Fife et al., 2009). Notably, we have shown that, in the RIP-mOva model of T1D, blockade of PD-1 is not sufficient to restore antigen-mediated T cell arrest in advanced insulitis (Friedman et al., 2014). Thus, the mechanism by which T cells are suppressed in the islets during advanced stages of infiltration is unknown.

In order for T cells to form productive interactions with APCs displaying cognate antigen, T cells must navigate and search for antigen in a complex environment with a variety of cues using tightly controlled mechanisms. These cues, which can regulate the sensitivity to antigen, include ligands for adhesion molecules, the physical characteristics of the tissue, chemokines and cytokines, costimulatory and coinhibitory molecules, and presented antigen. The balance of these cues controls the efficiency versus the sensitivity of the T cell response to antigen by altering the number of APCs or targets searched and the thoroughness with which the APCs or targets are scanned (Krummel et al., 2016; Fowell and Kim, 2021).

Mononuclear phagocytes play a key role in the induction of T1D and its pathogenesis, but their role in modulating the immune response in the pancreatic islets during T1D is less clear. Resident and infiltrating mononuclear phagocytes are present in the islets during T1D, with the majority of both resident and infiltrating populations expressing CD11c (Jansen et al., 1994; Melli et al., 2009; Calderon et al., 2015; Friedman et al., 2014). Islet-resident mononuclear phagocytes are composed of macrophages and a minor population of CD103+ dendritic cells (Yin et al., 2012; Friedman et al., 2014), while islet-infiltrating mononuclear phagocytes are monocyte derived (Klementowicz et al., 2017) and contain macrophages and dendritic cells (Zakharov et al., 2020). CD11c-depletion studies indicate that CD11c+ populations have a role in driving disease at early stages of T1D, while at later stages CD11c+ cells may slow disease progression (Saxena et al., 2007). This dichotomy could be due to temporal changes in CD11c+ populations (Zakharov et al., 2020) and/or differing roles of CD11c+ cells in different tissue locations (i.e. pancreatic lymph node vs. islets). The requirement for BatF3-dependent dendritic cells for the initial break in T cell tolerance in the draining pancreatic lymph nodes explains the requirement for CD11c+ cells prior to insulitis (Ferris et al., 2014; Gagnerault et al., 2002; Turley et al., 2003). CD11c+ cells also dramatically enhance entry of lymphocytes into inflamed islets, providing another potential node of disease regulation (Sandor et al., 2019). Yet, following the initial break in immunological tolerance the role of mononuclear phagocytes within the islets has not been elucidated. The fact that T1D progression is not altered by the removal of the draining pancreatic lymph nodes following the establishment of insulitis (Gagnerault et al., 2002) suggests that the disease site in the islets may be the relevant site to interrogate. The function of the mononuclear phagocytes in the islets at different stages of islet infiltration is not clear.

MERTK, which is best known as a macrophage marker, is expressed on mononuclear phagocytes where its signaling plays an immunoregulatory role. MERTK is a member of the TAM family of receptor tyrosine kinases. TAM receptors and their ligands function in phagocytosis of apoptotic cells, tolerance induction, coagulation, and erythropoiesis (Lu and Lemke, 2001; Nagata et al., 1996; Nakano et al., 1997; Scott et al., 2001; Van Der Meer et al., 2014; Rothlin et al., 2015). MERTK-mediated apoptotic cell uptake results in immunoregulation of APCs and production of inflammation resolution mediators (Wallet et al., 2008; Cai et al., 2018). MERTK has also been implicated in T cell tolerance induction to self-antigens expressed by apoptotic beta cells in the NOD model (Wallet et al., 2008). It has been suggested that MERTK might be a checkpoint molecule that could be targeted to intervene in immune tolerance (Akalu et al., 2017; Cook et al., 2013), yet its role as such has not yet been clearly defined.

Having previously identified a transient regulation of T cells in T1D, we sought to identify the mediator of this immune regulation through interrogating the mechanism by which T cells are suppressed at the autoimmune disease site. Using *in vivo* imaging of the pancreatic islets, we found that CD11c+ cells prevent T cells from arresting and responding to their antigen via a MERTK-dependent mechanism. This result was mirrored in a solid tumor model, indicating the broad applicability of this mechanism. MERTK inhibition or deficiency drove decreased T cell motility leading to enhanced T cell scanning for cognate antigen and increased antigen-dependent sustained interactions of T cells with CD11c+ antigen-presenting cells. This led to a very rapid increase in antigen experience and lytic potential by islet-infiltrating T cells, which was restricted to the disease site. The increased T cell activation in the islets resulted in the rapid onset of type 1 diabetes in animals with pre-existing insulitis. Increased numbers of MERTK-expressing cells were also observed in the remaining intact islets of type 1 diabetes patients. These data indicate that MERTK signaling in mononuclear phagocytes is driving tissue protective T cell regulation specifically at the autoimmune and tumor disease sites.

## Results

### CD11c+ cells mediate the suppression of T cell arrest in the islets

We previously identified temporally controlled periods of T cell stimulation and suppression in the islets (Friedman et al., 2014; Lindsay et al., 2015). Advanced stages of islet infiltration were dominated by T cell regulation which was characterized by a loss of: antigen-mediated T cell arrest, pathogenic cytokine production, and interactions with CD11c^+^ cells (Friedman et al., 2014; Lindsay et al., 2015). Since dendritic cells (DCs), macrophages, and monocytes all have the potential to drive T cell tolerance, we hypothesized that a subset of CD11c+ cells is responsible for this T cell regulation in advanced infiltrated islets. To test this hypothesis, we generated mixed bone marrow (BM) chimeras using NOD.CD11c-DTR (diphtheria toxin receptor) BM with either fluorescent islet-antigen specific 8.3 CD8 T cells (Fig 1A-G, Movie S1) or fluorescent polyclonal NOD T cells (Fig 1H-K). 5-10% fluorescently-labeled T cells were used to enable tracking of individual T cells. The chimeric mice were treated with two doses of diphtheria toxin (DT) to deplete the CD11c+ cells or vehicle (PBS) as a control (Sandor et al., 2019). Forty-eight hours after the initial treatment, islets were isolated for analysis (Fig 1A,H). We confirmed that CD11c depletion was effective in the islets (Fig S1A). The population depleted included most of the islet mononuclear phagocyte compartment, including CD11b+, XCR1+, and MERTK+ populations (Fig S1B).

**Figure 1:**
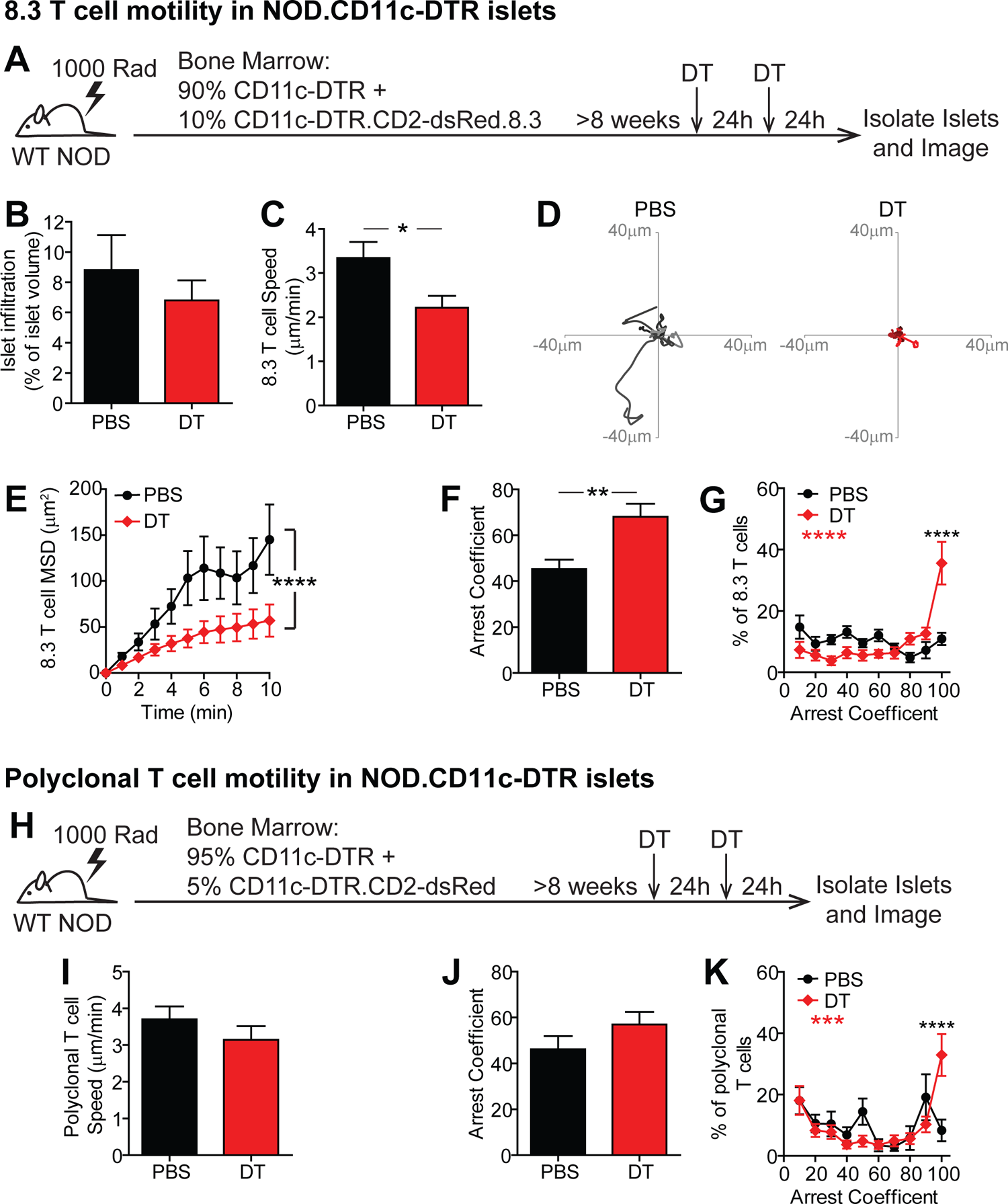
CD11c+ cells in the islets prevent T cell arrest. NOD.CD11c-DTR bone marrow (BM) chimeras containing **(A-G)** 90% NOD.CD11c-DTR BM +10% NOD.CD11c-DTR.8.3.CD2-dsRed BM or **(H-K)** 95% CD11c-DTR + 5% CD11c-DTR.CD2-dsRed were treated twice with PBS (Control) or DT (CD11c depleted). **(A,H)** Schematic of experimental setup to analyze T cell motility in explanted islets by 2-photon microscopy. **(B)** Average level of islet infiltration. **(C,I)** Average T cell crawling speed per islet. **(D)** Example of representative 15 min tracks of motion of 15 randomly selected T cells from individual islets. **(E)** Mean squared displacement (MSD) over time indicates the T cells’ ability to translocate away from their place of origin. Averaged per islet. **(F,G,J,K)** Arrest coefficient: % of time a T cell is moving <1.5 µm/min. Averaged per islet. **(A-K)** Data pooled from 3 independent experiments. Error bars represent SEM. **(A-G)** n = 10 islets containing 647 analyzed 8.3 T cells from PBS treated mice and n = 11 islets containing 371 analyzed 8.3 T cells from DT treated mice. **(H-K)** n = 13 islets containing 156 analyzed polyclonal T cells from PBS treated mice and n = 15 islets containing 329 analyzed polyclonal T cells from DT treated mice. **(B,C,F,I,J)** Statistics: Students T test. **(E,G,K)** Statistics: Two-way ANOVA with Bonferroni’s multiple comparison test, Treatment effect (black statistic), Interaction effect (red statistic). **(B-K)** *p<0.05, ***p<0.001 ****p<0.0001.

The ex vivo method of explanted islet imaging recapitulates T cell behavior in islets within the intact pancreas of live animals (Lindsay et al., 2015). The infiltration state of each islet analyzed was quantified to ensure that equivalent islets were analyzed in the intact and CD11c-depleted conditions (Fig 1B), since T cell behavior varies with the level of T cell infiltration (Lindsay et al., 2015; Friedman et al., 2014).

The depletion of CD11c+ cells resulted in significantly decreased islet antigen specific 8.3 CD8 T cell crawling speed within the islets (Fig 1C-D, Movie S1). The reduced crawling speed resulted in significantly reduced T cell translocation from the point of origin, as indicated by the mean squared displacement over time (Fig 1E). Furthermore, the 8.3 T cells in the islets showed greater arrest as indicated by an increased arrest coefficient (% of time a T cells spends crawling <1.5μm/min) (Fig 1F-G). There was a highly significant increase in the frequency of fully arrested T cells (arrest coefficient of 100) (Fig 1G). Notably, we previously determined that T cell arrest in the islets is antigen dependent and is correlated with increases in both T cell-APC interactions and T cell effector functions in the islets (Lindsay et al., 2015; Friedman et al., 2014), suggesting that the depletion of islet CD11c+ cells may be promoting T cell restimulation.

To determine whether CD11c+ cells have a similar effect on the motility and arrest of endogenous polyclonal CD4 and CD8 T cells in the islets, we depleted CD11c+ cells in mice with fluorescently-labeled polyclonal T cells (Fig 1H). The average T cell crawling speed (Fig 1I) and average T cell arrest coefficient (Fig 1J) were not significantly different between the CD11c-depleted and intact mice. This suggests that the CD11c+ cell dependent change in islet T cell motility is dependent on islet-antigen specificity. However, there was a highly significant increase in fully arrested T cells (arrest coefficient of 100) following CD11c+ cell depletion (Fig 1K), suggesting that a subset of islet-antigen specific polyclonal T cells is responding to CD11c+ cell depletion.

These data suggest that in the absence of suppressive CD11c+ cells, CD8 T cells are able to respond to antigen presented by other cells in the islets, such as beta cells, CD11c^-^ macrophages, or B cells, resulting in T cell arrest. The reduced effect on polyclonal T cells at a population level is likely due to the presence of non-islet antigen specific T cells in the islets and also the inability of CD4 T cells to recognize antigen on MHC class II^-^ beta cells. These data indicate that CD11c+ cells play a dominant role in preventing T cell arrest on APCs or beta cells in the islets.

DCs can have the potential to be either stimulating or tolerogenic. We next sought to determine whether Zbtb46-expressing classical DCs within the islets were responsible for regulating T cell arrest. Due to the availability of Zbtb46-DTR mice on the C57BL/6 background, we chose to use the RIP-mOva system in which a membrane bound form of ovalbumin (mOva) is expressed in the beta cells under the rat insulin promoter (RIP). We generated bone marrow chimeras using B6.Zbtb46-DTR (Meredith et al., 2012) bone marrow transferred into irradiated C57BL/6.RIP-mOva recipients (Fig S2A). In this system the transfer of ovalbumin-specific OT-I CD8 T cells induces T cell infiltration of the islets (Kurts et al., 1997). Our previous work has shown that islet-antigen specific OT-I T cells in RIP-mOva islets show the same dynamics of response as islet-antigen specific CD4 and CD8 T cells in the islets of NOD mice, including a period of T cell suppression in advanced infiltrated islets (Friedman et al., 2014). T cell infiltration of the islets was induced in the RIP-mOva.Zbtb46-DTR chimeric mice by transfer of naïve OT-I.Ubiquitin-GFP T cells. Three to five days following OT-I T cell transfer, the chimeric mice were treated with two doses of diphtheria toxin (DT) to deplete classical DCs or two doses of vehicle (PBS) as a control. Forty-eight hours after the initial dose, islets were isolated and imaged by 2 photon microscopy (Fig S2A). No differences were seen in the islet-antigen specific T cell crawling speed (Fig S2B) or arrest (Fig S2C) following classical DC depletion, despite equivalent infiltration of the islets analyzed (Fig S2D). Analysis of the islet mononuclear phagocyte populations following depletion in Zbtb46-DTR chimeras showed that XCR1+ cDC1s were significantly reduced, while CD11b+ and MERTK+ populations remained intact (Fig S2E). These data indicate that classical DCs are not required for the T cell regulation mediated by CD11c+ cells in the islets.

### MERTK is expressed on islet CD11c+ cells

Since CD11c+ cell-mediated T cell regulation was not dependent on classical DCs, we sought to understand the source of the regulation in macrophage and monocyte populations. Monocyte-derived CD11c+ cells infiltrate the islets with T cells (Melli et al., 2009; Jansen et al., 1994; Klementowicz et al., 2017). MERTK is expressed on macrophages and some monocytes in peripheral tissues (Guilliams et al., 2014; Tamoutounour et al., 2013), mediates immunoregulation of APCs, and has been implicated in T cell tolerance (Cabezón et al., 2015; Cook et al., 2013), including in the NOD model (Wallet et al., 2008).

We previously identified CX3CR1 levels as a marker to distinguish resident CD11c+ cells from infiltrating CD11c+ cells (Friedman et al., 2014). Here we show that in NOD islets, the numbers of islet-infiltrating CX3CR1^low^ CD11c+ cells directly correlated with total levels of islet infiltration (Fig 2A-C). MERTK was expressed on both islet-resident (CX3CR1^high^) and islet-infiltrating (CX3CR1^low^) CD11c+ cells in the NOD model (Fig 2A-B,D), and as previously observed, MERTK-expressing CD11c+ cells increased with increasing islet infiltration (Fig 2E) (Mohan et al., 2017). Thus, based on the increased frequency of MERTK-expressing CD11c+ cells in advanced infiltrated islets, we sought to determine if MERTK signaling was responsible for driving T cell regulation.

**Figure 2:**
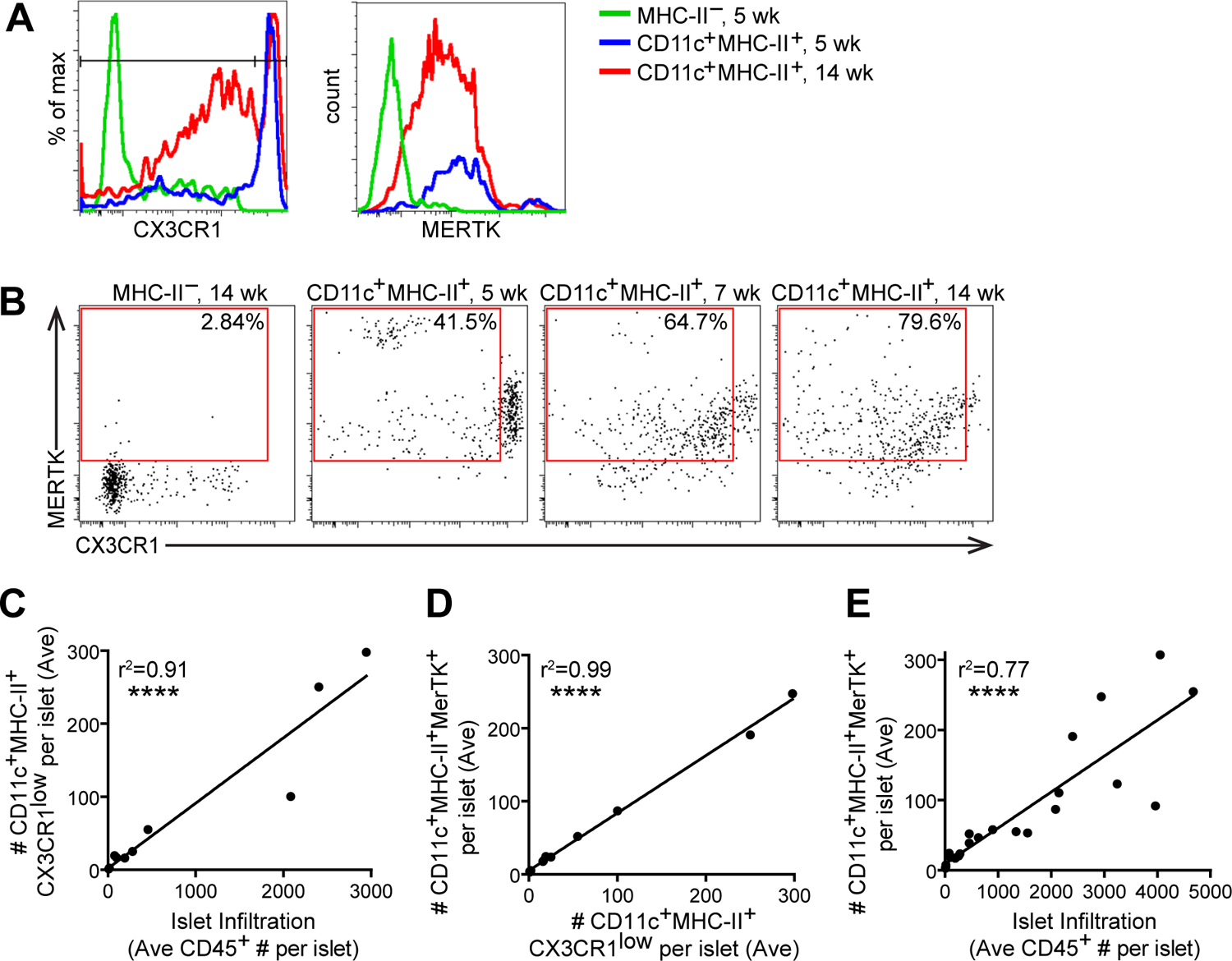
MERTK expressing cells increase with progression of islet infiltration. Islets from 4-18 week old NOD or NOD.CX3CR1-GFP female mice were isolated, digested and dissociated, stained, and analyzed by flow cytometry. **(A)** Example histograms of CX3CR1 gating in female NOD.CX3CR1-GFP islets. **(B)** Example plots of MERTK+CX3CR1^Low^ gating in female NOD islets, with % positive in the gate shown. **(C-E)** Each dot represents one mouse, as the # of cells in the islets normalized to the # of islets per mouse. **(C)** Quantification of CD11c^+^MHC-II^+^CX3CR^1ow^ (infiltrating APCs) versus the leukocyte infiltration in the islets. **(D)** Quantification of CD11c+MHC-II+MERTK^+^ cells versus CX3CR1^low^ CD11c+MHC-II+ infiltrating APCs in the islets. **(E)** MERTK+CD11c+MHC-II+ versus leukocyte infiltration in the islets. **(A-D)** n=11 mice from 2 independent experiments. **(E)** n=24 mice from 6 independent experiments. **(C,D,E)** Statistics calculated by Pearson correlation, ****p<0.0001.

### MERTK regulates islet T cell motility and arrest in a T cell-extrinsic manner

To test the hypothesis that MERTK signaling in mononuclear phagocytes promotes T cell regulation in infiltrated islets, we analyzed WT T cell motility in the islets of WT NOD versus NOD.MERTK^-/-^ mice. CD11c+ and CD11b+ mononuclear phagocyte frequency in the islets was not altered by MERTK deficiency (Fig S3A). However, NOD.MERTK^-/-^ mice have altered thymic selection which results in the absence of type 1 diabetes and strongly diminished spontaneous insulitis (Wallet et al., 2009). Thus, to induce islet infiltration, we transferred in vitro activated BDC-2.5 T cells into WT NOD and NOD.MERTK^-/-^ recipients. Three to six days later, fluorescently labeled activated 8.3 CD8 T cells and BDC-2.5 CD4 T cells were transferred into the same hosts. Twenty-four hours after fluorescent T cell transfer, the islets were isolated and analyzed by 2-photon microscopy (Fig 3A-B, Movie S2). WT and MERTK^-/-^ islets analyzed had equivalent infiltration (Fig 3C). Islet-antigen specific CD8 and CD4 T cell motility was significantly reduced in the islets of MERTK^-/-^ recipients compared to WT recipients (Fig 3D-E,I-J, Movie S2). The motility of the CD8 and CD4 T cells was also more confined in MERTK^-/-^ islets, as determined by the mean squared displacement (Fig 3F,K). This was accompanied by a significant increase in T cell arrest in MERTK^-/-^ islets (Fig 3G,L) with an increased frequency of fully arrested T cells in the islets of MERTK^-/-^ mice (Fig 3H,M). The increased T cell arrest observed in MERTK^-/-^ hosts suggests that MERTK signaling in MERTK-expressing CD11c+ cells drives the T cell-extrinsic regulation of T cell arrest in infiltrated islets.

**Figure 3:**
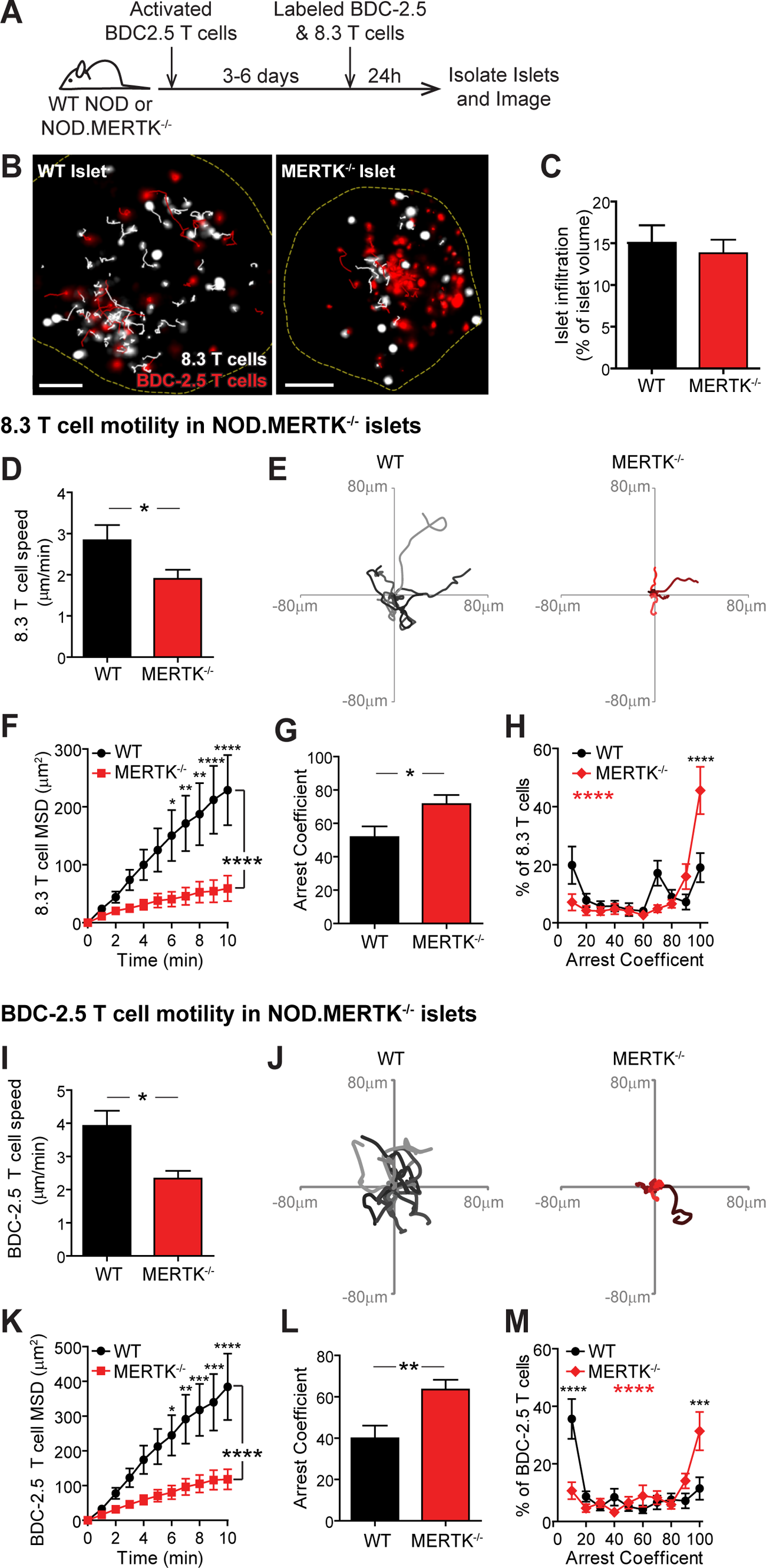
Islet antigen-specific T cells in MERTK-/- islets show decreased motility and increased T cell arrest. Activated BDC-2.5 CD4 T cells were transferred into NOD or NOD.MERTK-/- female mice to initiate islet infiltration. Fluorescently labeled activated 8.3 CD8 and BDC-2.5 CD4 T cells were then transferred into the same mice 3-6 days later. Twenty-four hours after fluorescent T cell transfer, islets were isolated and analyzed by 2-photon microscopy. **(A)** Schematic of experimental setup. **(B)** Representative islet images from WT NOD or NOD.MERTK-/- mice with 10 minute tracks of motion. Dashed line depicts islet border. 8.3 CD8 T cells are shown in white and BDC-2.5 CD4 T cells are shown in red. Scale bar = 50μm. **(C)** Average level of islet infiltration. **(D,I)** Average T cell crawling speed per islet. **(E,J)** Example of representative 15 min tracks of motion of 15 T cells from representative islets. **(F,K)** Mean squared displacement (MSD) over time. Averaged per islet. **(G,H,L,M)** Arrest coefficient: % of time that a cell crawls <1.5µm/min. Averaged per islet. **(C-M)** Data are pooled from 4-5 independent experiments. Error bars: SEM. **(D-H)** n= 22 WT islets containing 864 analyzed 8.3 T cells and n = 23 MERTK-/- islets containing 893 analyzed 8.3 T cells. **(I-M)** n= 25 WT islets containing 1069 analyzed BDC-2.5 T cells and n = 28 MERTK-/- islets containing 1496 analyzed BDC-2.5 T cells. **(C,D,G,I,L)** Statistics: Two-tailed Student t test. **(F,H,K,M)** Statistics: 2-way ANOVA with Bonferroni’s multiple comparison test, Treatment effect (black statistic), Interaction effect (red statistic). *p<0.05, **p<0.01, ***p<0.001 ****p<0.0001.

### MERTK signaling promotes T cell motility in infiltrated islets independent of cognate antigen

To determine whether MERTK signaling promotes T cell regulation in an antigen dependent or independent manner, we treated mice orally with UNC2025, a highly selective inhibitor of MERTK and Flt3 (Zhang et al., 2014). To ensure that UNC2025 was not reducing the CD11c+ or MERTK+ population in the islets via Flt3 inhibition, MERTK inhibition, or off-target effects, we analyzed the frequency of CD11c+ APCs and MERTK-expressing CD11c+ APCs in the islets following 4 weeks of continuous twice daily oral treatment with 30 mg/kg of UNC2025. There was no significant difference in these populations following 4 weeks of treatment (Fig S3B), thus we extrapolate that UNC2025 is not altering the frequency of the CD11c populations in the islets following short-term treatment.

To analyze islet antigen-specific T cell and polyclonal T cell motility, arrest, and displacement in the islets, we co-transferred activated, fluorescently-labeled, islet antigen-specific 8.3 CD8 T cells, islet-antigen specific BDC-2.5 CD4 T cells, and polyclonal NOD T cells into NOD.CD11c-YFP hosts. These recipient mice were then treated with UNC2025 or vehicle for 16 hours and then isolated islets were imaged by 2-photon microscopy (Fig 4A). The UNC2025 and vehicle treated islets analyzed were matched for level of infiltration (Fig 4B). Following UNC2025 treatment, there was a significant decrease in islet-antigen specific CD8 and CD4 T cell crawling speed as well as polyclonal NOD T cell crawling speed (Fig 4C). This was accompanied by increased T cell arrest in islet antigen specific and polyclonal T cells (Fig 4D-E). Islet-antigen specific CD8 and CD4 T cells showed a highly significant increase in fully arrested T cells following UNC2025 treatment while polyclonal T cells showed a modest increase in the frequency of fully arrested T cells following MERTK inhibition (Fig 4E). The movement of the islet antigen specific and polyclonal T cells was more confined following UNC2025 treatment compared to control saline treatment, as determined by the mean squared displacement over time (Fig 4F). Notably, mouse T cells do not express MERTK, thus MERTK-dependent T cell changes are extrinsically mediated by MERTK-expressing CD11c+ cells in the islets as supported by our findings in MERTK-/- mice (Fig 3). These data indicate that MERTK inhibition by UNC2025 treatment alters T cell motility in an antigen independent manner. These findings led to the question of whether this MERTK-mediated mechanism of T cell regulation was relevant to diverse disease settings.

**Figure 4:**
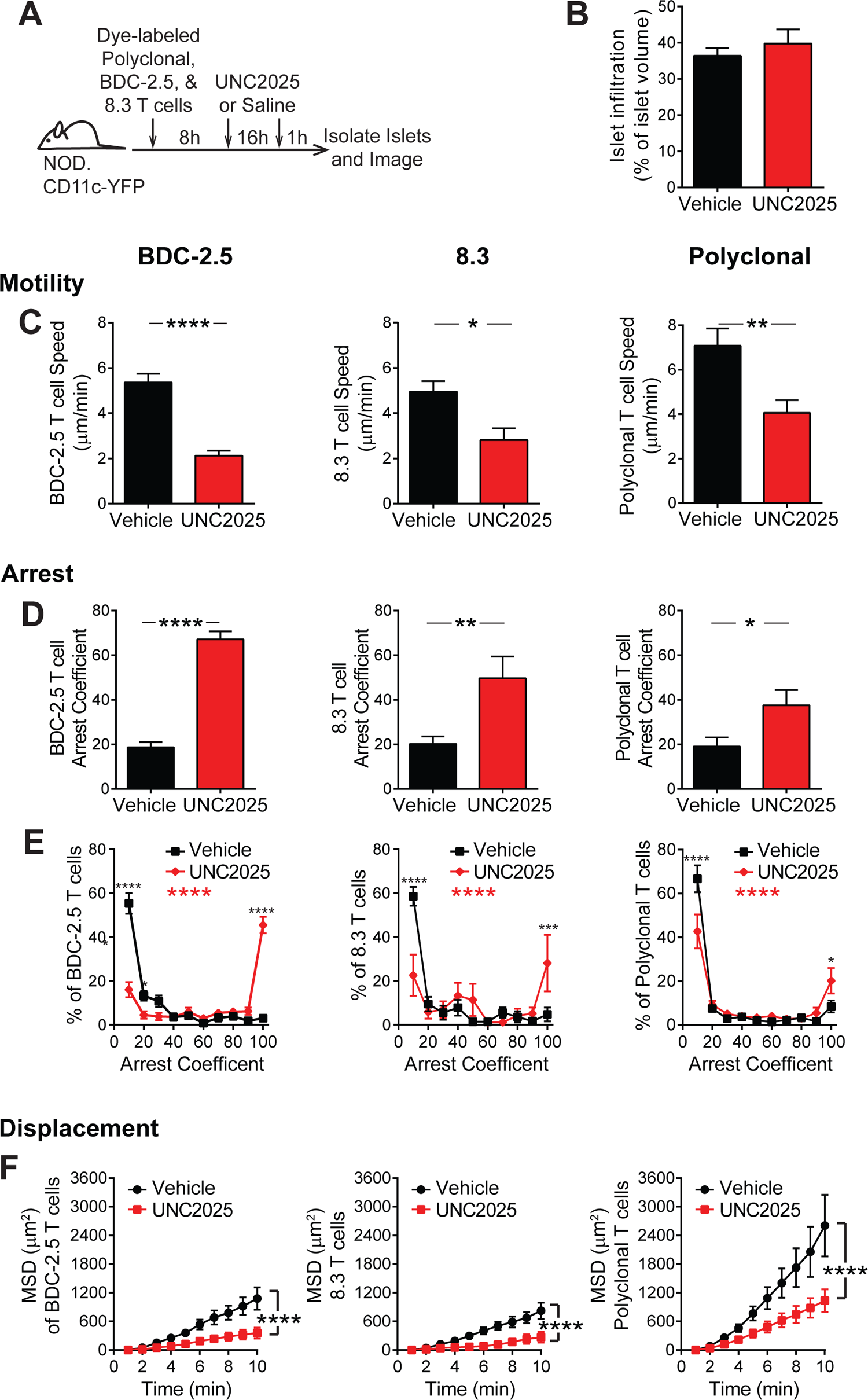
MERTK signaling drives antigen-independent suppression of T cell arrest in the islets. Activated 8.3 CD8 T cells, BDC-2.5 CD4 T cells, and Polyclonal T cells were fluorescently labeled and co-transferred into female NOD.CD11c-YFP mice. Mice were treated twice daily with saline vehicle or 30 mg/kg UNC2025, a small molecule inhibitor of MERTK and Flt3. 17h following the initial treatment, islets were isolated and analyzed by 2-photon microscopy. **(A)** Schematic of experimental setup. **(B)** Average level of islet infiltration. **(C)** Average T cell crawling speed per islet. **(D-E)** Arrest coefficient: % of time that a cell crawls <1.5µm/min. Averaged per islet. **(F)** Mean squared displacement (MSD) over time indicates the T cells’ ability to translocate away from their place of origin. Averaged per islet. **(A-F)** n= 17 islets from Vehicle treated mice, containing 590 analyzed BDC-2.5 T cells, 182 analyzed 8.3 T cells, and 497 analyzed Polyclonal T cells. n = 8-11 islets from UNC2025 treated mice, containing 481 analyzed BDC-2.5 T cells, 49 analyzed 8.3 T cells, and 671 analyzed Polyclonal T cells. Data were derived from 2-3 independent experiments. **(B-D)** Statistics: Two-tailed Student t test. Error bars: SEM. **(E-F)** Statistics: 2-way ANOVA with Bonferroni’s multiple comparison test, Treatment effect (black statistic), Interaction effect (red statistic), *p<0.05, **p<0.01, ***p<0.001 ****p<0.0001. Error bars: SEM.

### MERTK regulates T cell arrest in a solid tumor

Our findings indicate that MERTK signaling in mononuclear phagocytes regulates the T cell response in autoimmunity. Many mechanisms of immune regulation are shared between autoimmune and tumor settings, which led us to ask if regulation by MERTK signaling was altering the tumor-specific T cell response. TAM receptor family member mediated regulation of the T cell response within the tumor is an active area of clinical and pre-clinical research (Myers et al., 2019; Wu et al., 2018a; Zhou et al., 2020; Maimon et al., 2021). MERTK specific inhibition has been shown to suppress tumor growth in murine solid tumor models (Sinik et al., 2018; Wu et al., 2018b; Cook et al., 2013; Zhou et al., 2020). In some solid tumors, MERTK acts through modulation of the local type I IFN response (Zhou et al., 2020). Additionally, suppression of TAM receptor signaling has been shown to increase the response to anti-PD-1 therapy in murine tumor models, suggesting that it is mediated by a mechanism other than PD-1 signaling (Kasikara et al., 2019; Zhou et al., 2020). As a result, we sought to determine if a similar inhibition of T cell motility within solid tumors was observed with MERTK deficiency or following MERTK inhibition.

To determine whether MERTK regulates T cell motility in a solid tumor setting, we analyzed T cell motility and arrest in B78ChOva tumors. We chose the B78ChOva tumor model because it has a high frequency of MERTK-expressing tumor associated macrophages and expresses ovalbumin (Broz et al., 2014). B78ChOva tumor cells were implanted subcutaneously in C57Bl/6 mice and allowed to grow for 4-6 days. Activated tumor-specific OT-I T cells were then transferred. Following T cell trafficking to the tumor site, mice were treated with either saline or UNC2025 for 18h. Tumors were then excised and imaged by 2 photon microscopy (Fig 5A). To ensure that the effect was MERTK specific and to show that the effect was specific to the host cells recruited to the tumor, these experiments were repeated comparing WT and MERTK^-/-^ hosts.

**Figure 5:**
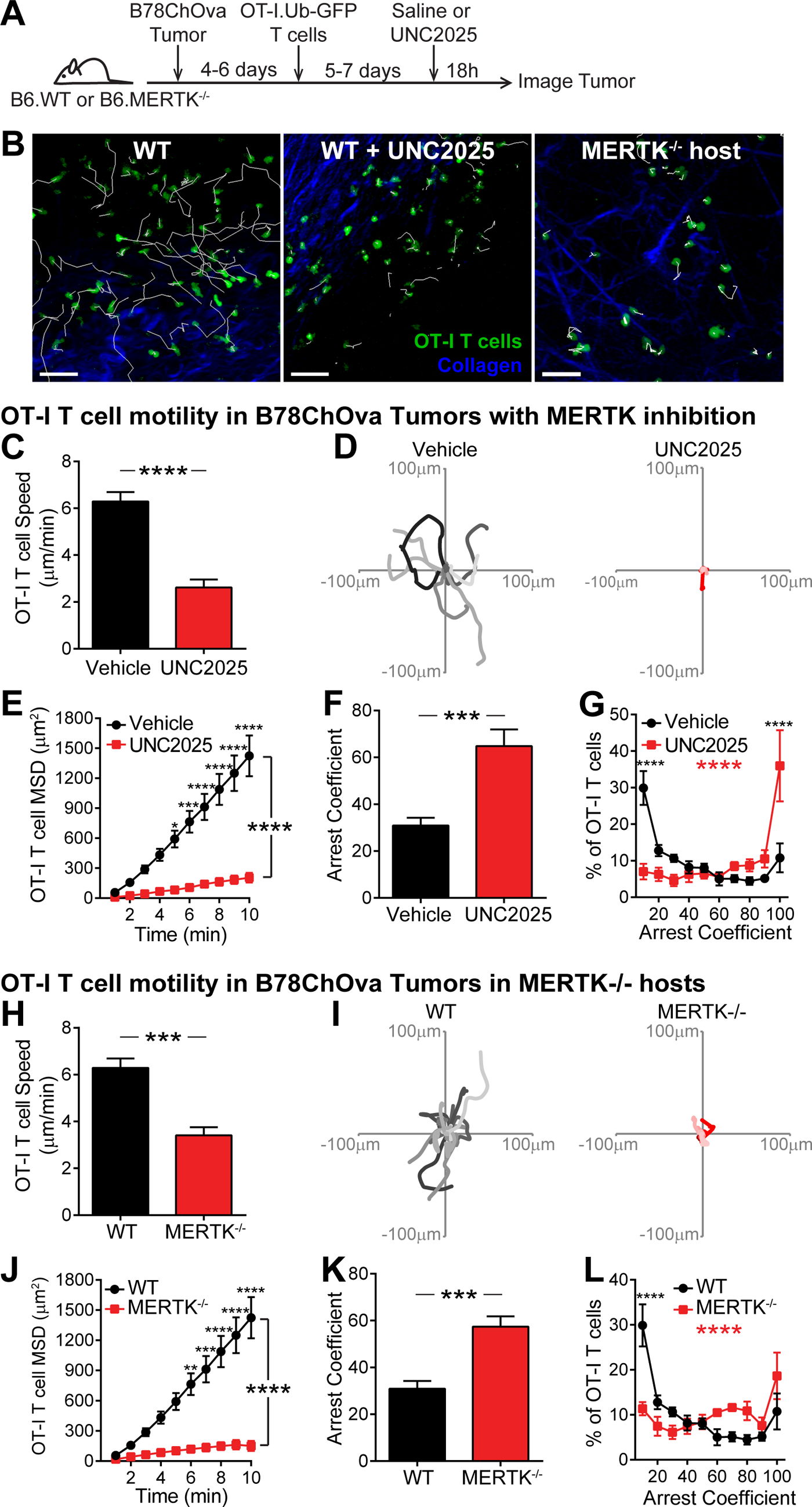
MERTK signaling in tumor associated host cells prevents tumor-specific T cell arrest in a solid tumor. B78ChOva tumor cells were implanted in C57BL/6 or C57BL/6.MERTK-/- mice. Activated OT-I.Ub-GFP CD8 T cells were then transferred into tumor bearing mice mice 4-6 days later. Five to seven days after fluorescent T cell transfer, mice were treated with UNC2025 or vehicle. 18h after treatment, tumors were excised and analyzed by 2-photon microscopy. **(A)** Schematic of experimental setup. **(B)** Representative images of OT-I T cell (green) motility on the tumor surface with 10 minute tracks of motion. Collagen (blue) delineates the tumor capsule. Scale bar = 50µm. **(C,H)** Average T cell crawling speed in the tumor. Averaged per tumor. **(D,I)** Example representative 15 min tracks of motion of OT-I T cells in representative tumors. **(E,J)** Mean squared displacement (MSD) over time. Averaged per tumor. **(F,G,K,L)** Arrest coefficient: % of time that a cell crawls <2µm/min. Averaged per tumor. **(B-L)** Data are pooled from 3-6 independent experiments. Error bars: SEM. n = 10 tumors from WT hosts, containing 2230 analyzed OT-I T cells. n = 6 tumors from UNC2025 treated mice, containing 1130 analyzed OT-I T cells. n = 5 tumors from MERTK-/- hosts, containing 641 analyzed OT-I T cells. **(C,F,H,K)** Statistics: Two-tailed Student t test. **(E,G,J,L)** Statistics: 2-way ANOVA with Bonferroni’s multiple comparison test, Treatment effect (black statistic), Interaction effect (red statistic), *p<0.05, **p<0.01, ***p<0.001 ****p<0.0001.

The tumor specific OT-I T cells showed a dramatic change in motility following MERTK inhibition (Fig 5B-G) or in MERTK^-/-^ hosts (Fig 5H-L). As in advanced infiltrated islets, OT-I T cells on the surface of the tumor were highly motile despite an abundance of local cognate antigen. However, when MERTK signaling was inhibited or deficient, the T cells increased arrest (Fig 5, Movie S3). Both the T cells and tumor cells that were transferred into MERTK-/- hosts were WT in origin, and thus contained an intact MERTK gene. This indicated that host derived cells, likely MERTK-expressing tumor associated macrophages, were responsible for the changes in T cell behavior. We next asked whether the T cell arrest promoted by MERTK inhibition was in part associated with increased antigenic stimulation.

### MERTK signaling regulates antigen-dependent T cell-APC interactions in the islets

While MERTK inhibition or deficiency modulated T cell motility and arrest in the islets in an antigen-independent manner, the higher speed and displacement of polyclonal T cells compared to islet-antigen specific T cells (Fig 4) raised the question of whether cognate antigen also played a role in this regulation. Previous work has suggested that MERTK-dependent tolerance is mediated by modulation of costimulation, including CD40, CD80, and CD86 (Wallet et al., 2008). We find that these costimulatory ligands and PD-L1 are not modulated on CD11c+ cells in the islets by inhibition of MERTK (Fig S4), suggesting an alternative mechanism of T cell regulation. Thus, we next explored if prevention of interactions between T cells and APCs might be a possible mechanism of MERTK mediated regulation.

To determine whether antigen-dependent T cell-APC interactions in the islets were increased as a result of MERTK inhibition, we analyzed 8.3 CD8 T cell, BDC-2.5 CD4 T cell, and polyclonal T cell interactions with CD11c+ cells following 16h of UNC2025 treatment (Fig 6A). In the lymph nodes, brief T cell-APC interactions have been shown to be tolerogenic; whereas sustained T cell-APC interactions have been shown to be activating (Jacobelli et al., 2013). Thus, we analyzed the duration of T cell-CD11c+ cell interactions and the frequency of sustained interactions longer than 10 minutes in duration. The frequency of islet-antigen specific CD4 and CD8 T cells participating in sustained interactions with CD11c+ cells was significantly increased in BDC-2.5 CD4 T cells, while polyclonal T cells showed no change (Fig 6C-D, Movie S4). Islet-antigen specific BDC-2.5 CD4 T cells also showed a significant increase in the duration of interaction with CD11c+ APCs in the islets following UNC2025 treatment, while polyclonal T cells showed no change (Fig 6E, Movie S4). These data suggest that while MERTK signaling alters T cell motility in an antigen-independent manner, it controls T cell interactions with APCs in an antigen dependent manner, by diminishing the efficiency of T cell to scanning for cognate antigen.

**Figure 6:**
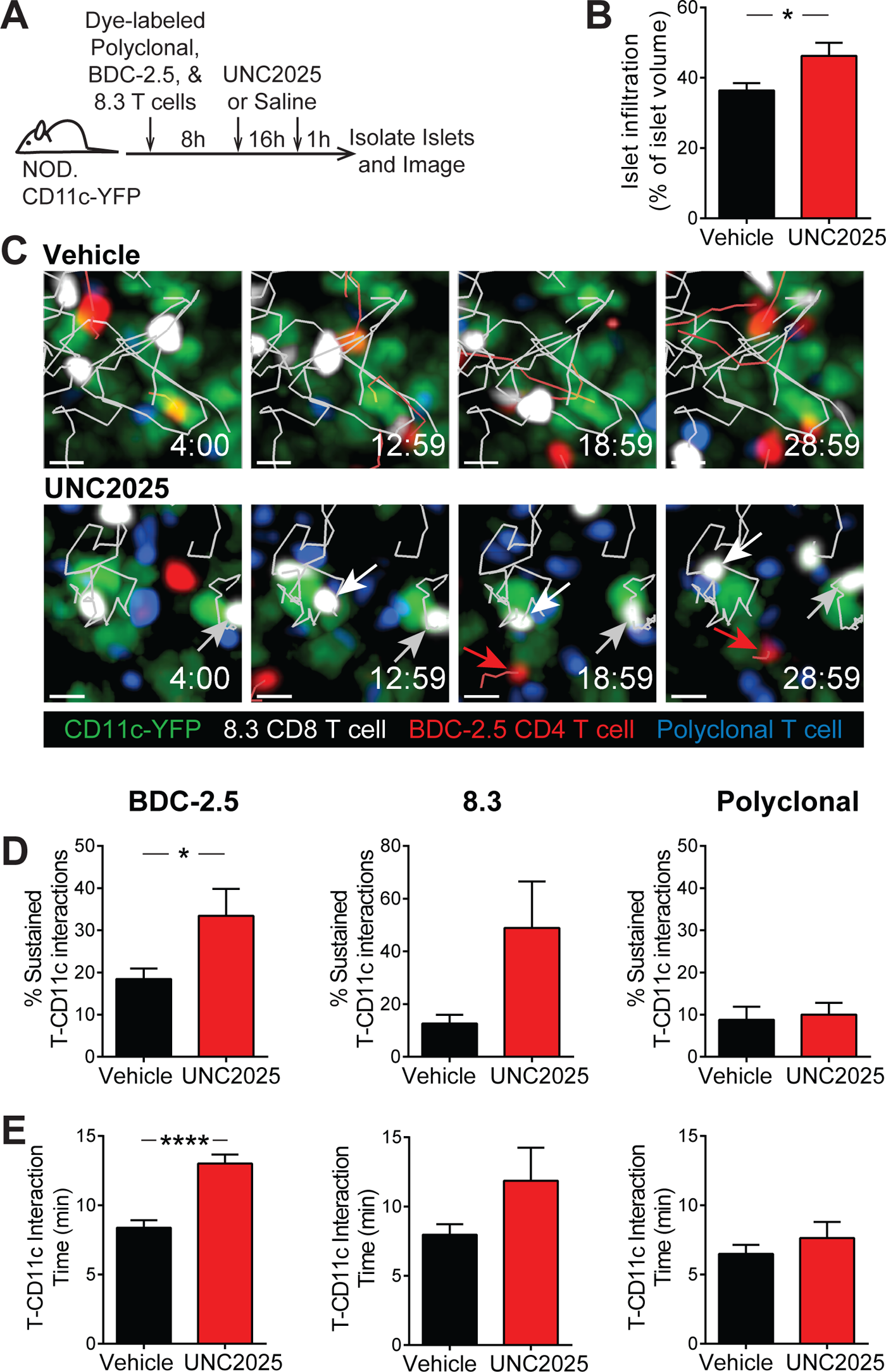
MERTK signaling drives suppression of antigen-dependent T cell-APC interactions in the islets. Activated 8.3 CD8 T cells (white), BDC-2.5 CD4 T cells (red), and Polyclonal T cells (blue) were fluorescently labeled and co-transferred into female NOD.CD11c-YFP mice (green). Mice were treated twice daily with saline vehicle or 30 mg/kg UNC2025. 17h following the initial treatment, islets were isolated and analyzed by 2-photon microscopy. **(A)** Schematic of experimental setup. **(B)** Average level of islet infiltration. **(C)** Representative islet images from Vehicle or UNC2025 treated mice with 5 minute tracks of motion. White and gray arrows point to sustained interactions between 8.3 T cells and CD11c cells while **(D)** % of T cells participating in sustained interactions (≥10 min) with CD11c+ cells. Scale bar = 10μm. **(E)** Average duration of interaction between T cells and CD11c+ cells. **(A-E)** n= 16-17 islets from Vehicle treated mice, containing 575 analyzed BDC-2.5 T cells, 168 analyzed 8.3 T cells, and 470 analyzed Polyclonal T cells. n = 8-9 islets from UNC2025 treated mice, containing 356 analyzed BDC-2.5 T cells, 60 analyzed 8.3 T cells, and 907 analyzed Polyclonal T cells. Data were derived from 2-3 independent experiments. **(B,D,E)** Statistics: Two-tailed Student t test, *p<0.05, ****p<0.0001. Error bars: SEM.

### T cell antigen recognition and effector function are increased following MERTK inhibition

To determine how MERTK-mediated regulation of T cell-APC interactions affected antigen recognition and activation by T cells in the islets and lymph nodes, we analyzed expression of CD44 and CD62L by flow cytometry following 17h of UNC2025 treatment of NOD mice. Polyclonal CD8 and CD4 T cells in the islets showed a rapid and significant increase in antigen experience (CD44^high^CD62L^-^) following MERTK inhibition (Fig 7A, Fig S5), suggesting that the increased arrest and T cell-APC interactions seen with MERTK inhibition are driving increased antigen recognition and TCR signaling in the islets. Notably, there was no significant difference in antigen experience in the draining pancreatic lymph nodes or non-draining inguinal lymph nodes (Fig 7A, Fig S5). These data suggest that MERTK signaling is functioning to suppress the T cell response by preventing antigen recognition specifically at the disease site in the islets.

**Figure 7:**
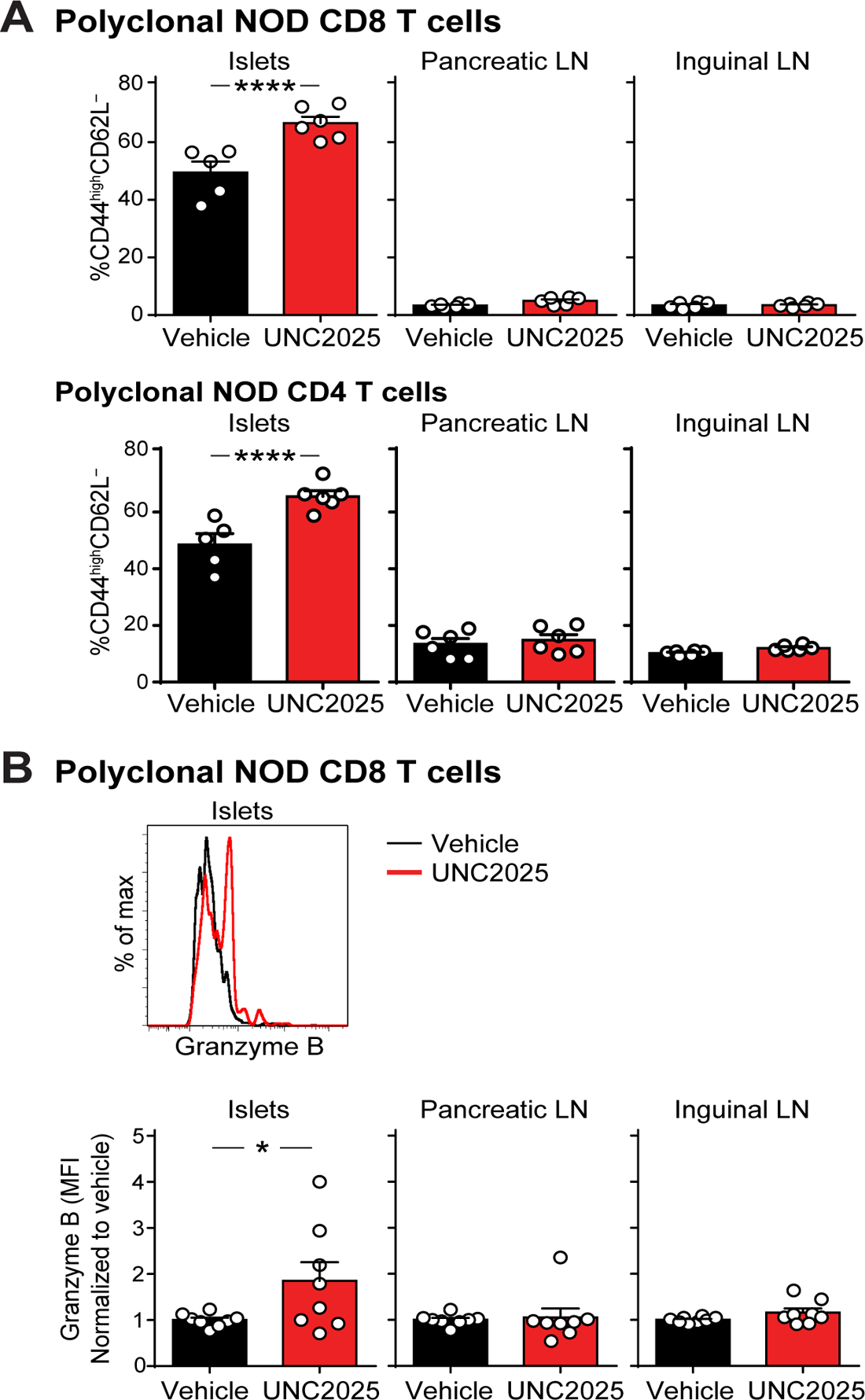
T cell antigen experience and effector function increases specifically at the disease site in the islets following MERTK inhibition. (A-B) NOD female mice were treated with UNC2025 or vehicle. Islets and lymph nodes were isolated 17h post-treatment initiation, digested, stained and analyzed by flow cytometry. **(A)** Quantification of antigen experience by CD44/CD62L expression. Gated on CD45^+^CD90.2^+^CD8^+^ or CD45^+^CD90.2^+^CD4^+^. n = 6 mice per group from 2 independent experiments. Statistics: 2-way ANOVA with Bonferroni post-test. Error bars: SEM. **(B)** Quantification of effector function by granzyme B expression. Mice were treated in vivo with Brefeldin A for 4 hours prior to harvest to block secretion. Cells were gated on CD45+CD8+Granzyme B+. Granzyme B expression was measured by normalized MFI. n = 8 mice per group from 4 independent experiments. Error bars: SEM. Statistics performed on the non-normalized MFI values using a Mann-Whitney test. *p<0.05, ****p<0.0001.

To determine whether effector function was also increased by MERTK inhibition, we analyzed expression of granzyme B in CD8 T cells by flow cytometry following 17-24h of UNC2025 treatment of NOD mice. Granzyme B expression serves as an indicator of CD8 T cell activation and lytic potential. Polyclonal CD8 T cells in the islets showed a rapid and significant increase in the amount of granzyme B following MERTK inhibition (Fig 7B). On the other hand, IFN-γ production was not modulated with MERTK inhibition (data not shown). As seen with antigen experience, there was no significant difference in granzyme B expression in the draining pancreatic lymph nodes or non-draining inguinal lymph nodes (Fig 7B), further indicating that MERTK signaling is acting specifically at the disease site. Notably, only half of the mice showed an increase in CD8 T cell granzyme B, indicating that while the T cells in the islets of all mice show increases in antigen experience, other mechanisms of tolerance may control effector function.

### Inhibition of MERTK signaling drives rapid type 1 diabetes induction in mice with existing islet infiltration

To determine how inhibition of MERTK affects type 1 diabetes progression, we treated NOD mice with UNC2025 or saline vehicle for 17h and analyzed changes in islet infiltration by quantification of histological staining. The data show a rapid increase in the frequency and degree of islet infiltration after only 17h of MERTK inhibition (Fig 8A). We also treated mice twice daily with UNC2025 or vehicle for 14 days and monitored blood glucose levels for diabetes. We chose ≥20 week old female NOD mice because we reasoned that this cohort would have a high proportion of infiltrated islets, where MERTK signaling might be driving T cell regulation. Additionally, we treated WT female C57Bl/6 mice with UNC2025 as an additional control to ensure that UNC2025 did not have direct beta cell toxicity. MERTK inhibition by UNC2025 treatment rapidly induced diabetes in a subset of the NOD mice (Fig 8B). The diabetes incidence was similar to the incidence of enhanced effector function following MERTK inhibition, suggesting that additional tolerance mechanisms within the islets may be protecting some of the mice. The rapid nature of disease onset suggests that MERTK is playing a critical role in regulating T cell pathogenesis in the islets to prevent rapid disease progression during type 1 diabetes. This led us to the question of whether MERTK also functions in human type 1 diabetes to protect islets from destruction.

**Figure 8:**
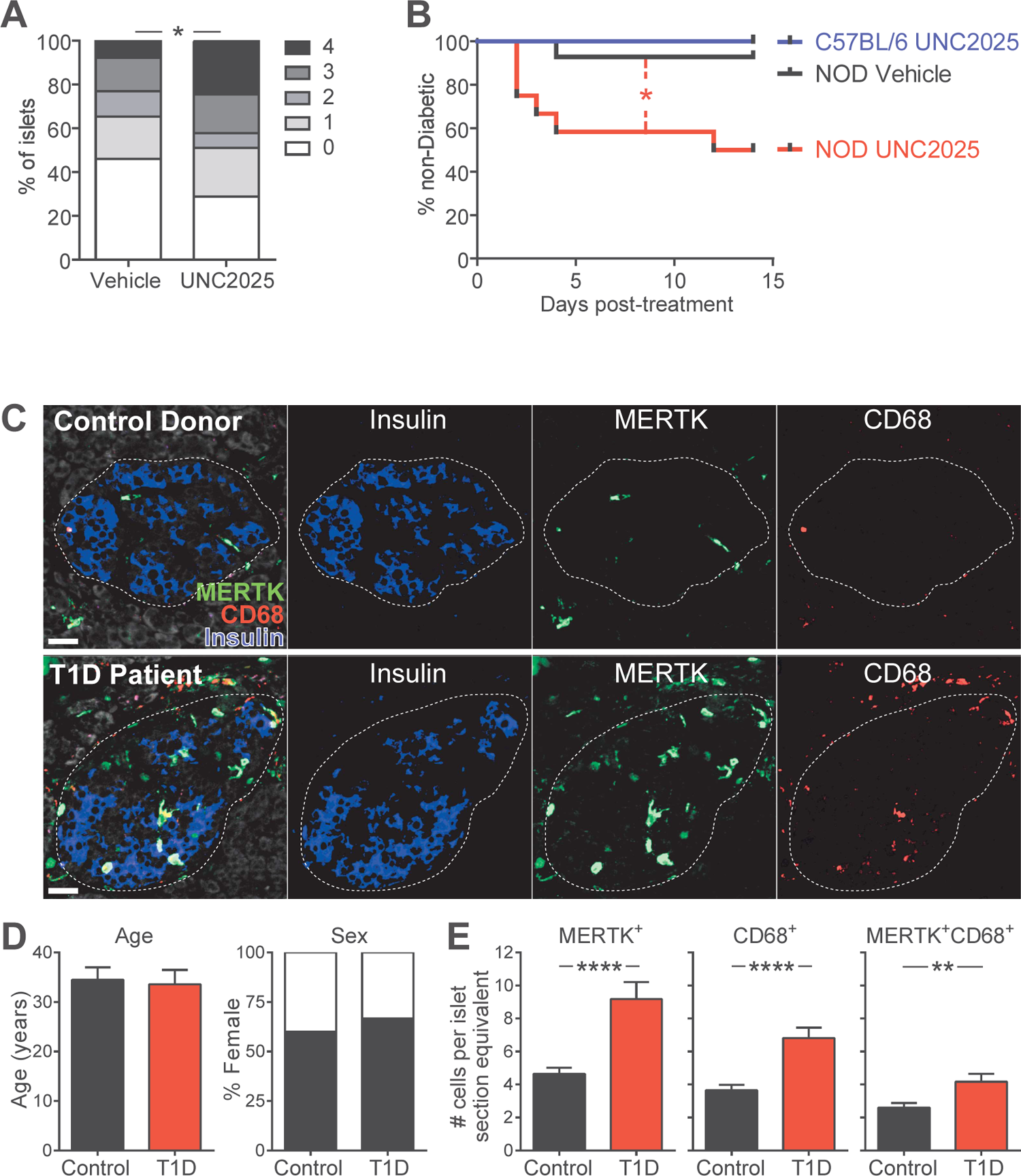
MERTK signaling prevents rapid T1D progression in mice and MERTK-expressing cells are increased in the remaining insulin-containing islets of T1D patients. (A-B) NOD female mice over 20 weeks of age or WT C57BL/6 female mice were treated orally twice daily with 30mg/kg UNC2025 or saline Vehicle twice daily for **(A)** 16 hours or **(B)** 14 days. **(A)** Quantification of islet infiltration following MERTK inhibition. Whole pancreata were isolated and fixed prior to sectioning and staining. Sections were scored blindly for islet infiltration using the standard insulitis scale of 0 = uninfiltrated islet to 4 = completely destroyed islet. n=45 islets per condition from 5-6 mice per group from 2 independent experiments. Statistics: Student’s T test of islet scores. **(B)** Diabetes incidence with MERTK inhibition. Mice were monitored daily for blood glucose values. Mice with two consecutive blood glucose readings of >250 mg/dL were considered to be diabetic. n = 6-14 mice per group from 4 independent experiments. Statistics: Mantel-Cox Test. **(C-E)** Immunofluorescent staining of pancreas sections from type 1 diabetes patient and control donors. **(C)** Representative images of MERTK staining (green), CD68 staining (red), insulin staining (blue), and tissue autofluorescence (white). The dotted line indicates the islet border. Scale bar = 30μm. **(D)** Donor information. **(E)** Blinded quantification of MERTK-expressing cell number per islet section. n= 118 islets from 9 T1D patients and 131 islets from 10 controls. Error bars = SEM. Statistics: Two-tailed Student t test. *p<0.05, **p<0.01, ****p<0.0001.

### Remaining intact islets of type 1 diabetes patients have increased numbers of MERTK-expressing cells

We hypothesized that if MERTK protects islets during type 1 diabetes, we would see an increased number of MERTK-expressing cells in the remaining intact islets of type 1 diabetes patients. Thus, we used immunofluorescent staining of pancreatic sections from control and type 1 diabetic donors obtained from nPOD (network of pancreatic organ donors with diabetes) to test this hypothesis (Fig 8C-E). Control and type 1 diabetic donors had a similar mean age and sex distribution (Control: 23.3-44 years, mean 34.5 years; T1D: 20.3-45 years, mean 33.6 years) (Fig 8D, Table S1). The type 1 diabetes patients had long-standing disease of 10-20 years (Table S1). Insulin-containing islets from type 1 diabetes patients showed a significant increase in the number of MERTK-expressing cells in the islets when compared to the islets of controls (Fig 8C,E). Cell numbers per islet were normalized to the average area of all islets analyzed. Co-staining with the macrophage marker CD68 showed increased macrophages and MERTK-expressing macrophages in insulin containing islets of type 1 diabetes patients compared to control islets. Not all MERTK-expressing cells in the islets expressed CD68, suggesting that other mononuclear phagocyte subsets express MERTK in human islets (Fig 8C,E). These data support our hypothesis that MERTK-expressing mononuclear phagocytes could protect human islets from autoimmune destruction.

## Discussion

In this study, we sought to determine the mechanism by which autoreactive T cells are regulated at the tissue site of disease. Our results indicate that, following inflammation, CD11c-expressing cells act to prevent T cells from effectively searching for and responding to antigen by promoting rapid T cell scanning of APCs with low sensitivity antigen detection. This mechanism of T cell regulation is not dependent on classical dendritic cells. Instead, MERTK signaling in macrophages and/or monocytes suppressed local T cell stimulation by preventing T cells from effectively scanning for antigen and responding to antigen-bearing APCs in the islets during type 1 diabetes and at a solid tumor site. Inhibition of this MERTK-mediated mechanism of T cell regulation led to increased antigen experience and effector function by T cells at the autoimmune site, but not in draining lymph nodes. Ultimately, inhibition of MERTK-mediated immune regulation led to the acceleration of autoimmune pathology and disease progression.

Increased numbers of MERTK-expressing cells were also observed in the remaining intact islets of type 1 diabetes patients. Together, these data indicate that MERTK signaling in mononuclear phagocytes drives T cell regulation at the disease site in peripheral tissues. MERTK-mediated regulation is achieved by reducing the sensitivity of T cell scanning of the local antigenic landscape, thereby suppressing detection of and responsiveness to antigenic stimuli.

Our previous work indicated that T cell stimulation in the autoimmune lesion is controlled by modulating antigen-mediated T cell arrest during disease progression (Friedman et al., 2014; Lindsay et al., 2015). In type 1 diabetes progression, mild insulitis was characterized by antigen-mediated T cell arrest and restimulation, while advanced insulitis was characterized by the loss of antigen-mediated T cell arrest and antigenic stimulation despite locally presented cognate antigen (Friedman et al., 2014; Lindsay et al., 2015). These data suggest that tolerogenic pathways are functioning to prevent antigen-mediated T cell arrest. In a variety of systems, T cell tolerance induction has been associated with a lack of T cell arrest and with transient T cell-APC interactions (Jacobelli et al., 2013). For example, prevention of antigen-mediated T cell arrest as a means of tolerance induction has been established for the PD-1 pathway (Fife et al., 2009), where PD-1 signaling dephosphorylates CD28 to prevent co-stimulation (Hui et al., 2017). Notably, we have shown that blockade of PD-1 is not sufficient to restore antigen-mediated T cell arrest in advanced insulitis (Friedman et al., 2014), and that PD-L1 expression is not altered by MERTK inhibition (Fig S4). While MERTK signaling is known to play a role in immune regulation (Wallet et al., 2009; Scott et al., 2001; Van Der Meer et al., 2014; Cabezón et al., 2015), it has not previously been implicated in suppressing the T cell response by inhibiting effective scanning for antigen and antigen-mediated T cell arrest. Here we show that inhibition of or deficiency in MERTK leads to increased T cell arrest within T1D islets and at a solid tumor site.

T cell behavior in the islets during T1D is controlled by the islet milieu, which is dependent on the disease state of the individual islet (Friedman et al., 2014). Here, we show that the MERTK-expressing monuclear phagocytes are responsible for controling the T cell response by altering the islet envirnoment. The T cell extrinsic nature of this process is highlighted by the fact that mouse T cells do not express MERTK and that T cell behavior in the islets and tumor were altered in WT T cells within a MERTK-deficient environment. The evidence for MERTK-expressing mononuclear phagocytes driving this effect is particularly strong in the tumor model where both the T cells and tumor carried a WT MERTK gene, and only the host cells recruited to the tumor, including mononuclear phagocytes, were MERTK-deficient.

While MERTK-mediated regulation is T cell-extrinsic, there is a possible role for T cells in promoting MERTK signaling in the mononuclear phagocytes. Activated T cells can express the MERTK ligand Protein S, and the T cell derived Protein S can act to restrict the activation of DCs (CarreraSilva et al., 2013). Additionally, viable T cells can expose patches of phosphatidyl serine on their surface upon antigen recognition (Fischer et al., 2006), which could serve together with Protein S as a ligand on T cells to induce MERTK signaling. Thus, while MERTK-mediated regulation is T cell extrinsic, it is possible that T cells contribute to the induction of MERTK signaling. In contrast to mouse T cells, human T cells can express MERTK, which can function as a costimulatory molecule, making the players responsible for MERTK signaling in human settings more complex (Peeters et al., 2019). Further studies will be needed in human disease settings to determine the precise role of MERTK signaling on different cell types.

Polyclonal T cells showed reduced motility in the islets following MERTK inhibition, showing that the altered islet milieu induced by MERTK inhibition modifies T cell behavior in an antigen independent manner. Yet, increased T cell-APC interactions were dependent on antigen specificity, indicating that enhancing T cell scanning increased the T cell’s ability to detect local cognate antigen. Patterns of T cell scanning for cognate antigen in target tissues such as the pancreatic islets or tumor are controlled by a variety of different cues from the local environment, including ligands for adhesion molecules, physical characteristics of the tissue, chemokines and cytokines, and presented antigen. The balance of these cues alters the efficiency and sensitivity of the T cell search for antigen (Krummel et al., 2016; Fowell and Kim, 2021). Our data suggest that MERTK signaling in mononuclear phagocytes establishes a regulatory milieu that promotes a T cell search pattern that prioritizes high speed scanning over sensitivity to antigen. We believe that this represents one of the mechanisms by which MERTK signaling functions to reestablish homeostasis following an immune response. While mononuclear phagocytes in the islet environment utilize MERTK signaling to try to reestablish homeostasis, in T1D the failure of this pathway enables continued destruction of the islets. MERTK signaling in tumor associated mononuclear phagocytes enables the tumor to escape T cell surveillance by reducing T cell sensitivity to local antigen.

MERTK signaling suppresses the NFkB pathway (Sen et al., 2007; Zhang et al., 2017; Camenisch et al., 1999).NFkB target genes include the pro-inflammatory cytokines TNF-α, IL-1β, IL-6, and IL-12p40 (Liu et al., 2017),which can all be expressed by mononuclear phagocytes and can be suppressed by the MERTK signaling axis (Zhang et al., 2017; Wallet et al., 2008; Camenisch et al., 1999; Maimon et al., 2021).Given the role of the islet milieu in controlling T cell responsiveness in the islets, (Friedman et al., 2014), this leads us to hypothesize that MERTK signaling creates a regulatory milieu by modulating soluble mediators such as cytokines that alter T cell scanning by tuning T cell adhesion and responsiveness to antigen.

Many mechanisms work in concert to maintain and restore self-tolerance. This might be in part why MERTK inhibition led to increased effector function and disease progression in only half of the mice treated. It is likely that other mechanisms of tolerance were able to compensate for MERTK in some of the subjects, preventing the increased effector function and islet destruction. Despite preventing increased effector functions, other tolerance mechanisms were unable to prevent restoration of T cell arrest in both the islets and tumor. The asynchronous nature of disease in the NOD model might also play into this dichotomy since our data suggest that MERTK is functioning specifically within infiltrated islets. Thus, the percentage of uninfiltrated islets, which will not be destroyed upon MERTK inhibition, might determine whether or not MERTK inhibition results in overt diabetes. Co-inhibitory molecules such as CTLA-4, PD-1 and LAG-3 (Ansari et al., 2003; Keir et al., 2006; Bettini et al., 2011; Lühder et al., 1998; Luhder et al., 2000) as well as Tregs also provide layers of tolerance in the progression of type 1 diabetes (Tang et al., 2004a; Tarbell et al., 2007; Tonkin and Haskins, 2009). These regulatory mechanisms have roles in both the draining lymph node and at the disease site (Tang et al., 2004b; Bettini et al., 2011; Keir et al., 2006; Fife et al., 2009; Lühder et al., 1998; Luhder et al., 2000; Tarbell et al., 2007; Tonkin and Haskins, 2009). The role of MERTK in driving T cell tolerance following well established inflammation at the disease site could make it of particular importance when considering mechanisms that are critical in targeting tolerance of established disease at the site of pathology.

T1D has been proposed to be a relapsing remitting disease (von Herrath et al., 2007; van Megen et al., 2017), thus the temporal regulation of T cell pathogenesis in the islets could represent relapsing and remitting phases of disease within individual islets. Cycles of MERTK signaling initiation in the presence of β cell apoptosis followed by cessation once the autoimmune β cell destruction is controlled could control cycles of quiescent T cell tolerance followed by destructive re-initiation of T cell pathogenesis.

In this study, we identified MERTK signaling as a key mechanism in driving T cell regulation within the pancreatic islets during type 1 diabetes and at the tumor site. It is known that MERTK deficiency can also result in a systemic, lupus-like autoimmune phenotype (Scott et al., 2001). Furthermore, we would posit that this pathway is likely to inhibit antigen-mediated T cell arrest and stimulation in peripheral tissues under many inflammatory conditions including other types of tissue specific autoimmunity and in other solid tumors. In fact, MERTK expression in tumor infiltrating macrophages has been implicated in enabling tumor growth and metastasis in MMTV-PyVmT and B16:F10 tumor models (Cook et al., 2013). Antibody blockade of MERTK signaling was also recently shown to enhance T cell function in tumors by increasing type I IFN production by macrophages (Zhou et al., 2020). In future studies it will be important to understand the breadth of function of this MERTK-dependent tolerance pathway, why it fails in autoimmune settings, how it controls the sensitivity of T cell scanning for antigen, and how it inhibits T cell responsiveness to antigen.

## Materials and Methods

### Mice

NOD/ShiLtJ (Jackson 001976), NOD.8.3 (Jackson 005868), C57BL/6.RIP-mOva (Jackson 005431), C57BL/6.OT-I (Jackson 003831), C57BL/6.ZBTB46-DTR (Jackson 019506) and C57BL/6.Ubiquitin-GFP (Jackson 004353), C57BL/6.MERTK-/-(Jackson 011122) mice were obtained from The Jackson Laboratory and bred in-house. NOD.BDC-2.5 TCR transgenic (Katz et al., 1993),were a generous gift from the laboratory of Kathryn Haskins. NOD.CD11c-DTR (Saxena et al., 2007),mice were a generous gift from the laboratory of Jonathan Katz. NOD.CD2-dsRed mice were a generous gift from the laboratory of Qizhi Tang. NOD.MERTK-/- mice were a generous gift from the laboratory of Roland Tisch. NOD.CX3CR1-GFP and NOD.CD11c-mCherry mice were generated by backcrossing. NOD.CX3CR1-GFP and NOD.CD11c-mCherry were confirmed to be fully backcrossed to our in-house NOD mice utilizing speed congenic analysis performed by the Barbara Davis Center core facilities. All NOD transgenics and knockouts were backcrossed to NOD/ShiLtJ for at least 10 generations. All animal procedures were approved by the Institutional Animal Care and Use Committee at National Jewish Health and University of Colorado Anschutz Medical Campus.

### Islet isolation

Islets were isolated as previously described (Melli et al., 2009; Friedman et al., 2014).Briefly mice were anesthetized with Ketamine/Xylazine prior to cervical dislocation. The pancreas was inflated via the common bile duct by adding ∼3 ml of 0.8mg/ml Collagenase P (Roche) and 10µg/ml Dnase I (Roche) or CIzyme RI (Vitacyte) in HBSS (Cellgro). Following inflation, the pancreas was removed and incubated at 37°C for 10-11 min for collagenase P digestion or 17 min for CIzyme RI digestion and the islets were isolated by density centrifugation. The islets were handpicked under a dissecting microscope. For experiments in which islets were analyzed using flow cytometry single cell suspensions were generated by digestion with 0.4 Wunsch Units/ml Collagenase D (Roche) and 250 µg/ml DNAse I (Roche) in HBSS (Cellgro) with 10% FBS (Hyclone) at 37°C for 30 min. Following initial digestion islets were then incubated in Cell Dissociation Buffer (Sigma) at 37°C for an additional 30 min.

### Tumor implantation and excision

1×10^5^ B78ChOva tumor cells (generous gift of Max Krummel) were resuspended in PBS and mixed 1:1 with growth factor reduced Matrigel (Corning). Tumor cells were injected subcutaneously into each flank of WT or MERTK-/- C57BL/6 mice. At the time of harvest, tumors were carefully exposed by peeling the skin away from the peritoneum and excised with a border around the tumor. The skin associated with the tumor was affixed to a cover slip using VetBond (3M).

### 2-photon imaging of explanted islets and tumors

For 2 photon imaging, isolated islets were embedded in 3% low melting temperature agarose (Fisher) in DPBS. During imaging, islets and tumors were maintained at 35-37°C with flow of 95% O2/5% CO2 saturated RPMI (Gibco). Islets and tumors were imaged on an Olympus FV100MPE microscope or a Leica SP8 DIVE upright multi-photon microscope as previously described (Friedman et al., 2014; Lindsay et al., 2015). Excitation was performed at 810nm for islets and 910nm for tumors. Imaging fields were as described (McKee et al., 2013), xy planes of 509 µm by 509 µm with a resolution of 0.994 µm/pixel or xy planes of 592 µm by 592 µm with a resolution of 1.16 µm/pixel were acquired. Images of 27-60 xy planes with 3-um z-spacing were acquired every minute for 30 minutes. Four emission channels were used for data acquisition, blue (450- to 490-nm), green (500- to 550-nm), red (575- to 640-nm), and far-red (645- to 685-nm). In tumors collagen was imaged in the blue channel through second harmonic generation. The collagen was confirmed to be at the surface of the tumor by visualization of the mCherry tumor fluorescence at 810nm.

### Image Analysis

Image analysis was performed using Imaris (Bitplane) and MATLAB (Mathworks). Images were linearly unmixed, as previously described (Bullen et al., 2009). Islet infiltration levels were determined as previous described (Lindsay et al., 2015). Islets with similar levels of infiltration between groups were used for comparison. Cells in the islets that could be tracked for ≥5 min were used to obtain speed and average arrest coefficient (<1.5 µm/min cutoff for islets due to overall lower T cell speeds in the islets; <2 µm/min cutoff for tumors). Cells tracked for ≥10 min in the islets were used to determine mean squared displacement to ensure all cells analyzed could be included for all time points of the analysis. Interaction duration was calculated using a maximum distance between cells of 1 pixel (0.994 µm), with a minimum interaction of 2 timepoints to be defined as an interaction. Sustained interactions were defined as ≥10 min long interactions. Statistics were calculated with Prism software (GraphPad).

### Activation and dye labeling of BD2.5, 8.3, Polyclonal NOD, and OT-I T cells

Spleen and lymph node cells from female BDC-2.5 or 8.3 mice or male/female OT-I.Ub-GFP mice were stimulated in vitro with 1µg/ml of BDC-2.5 mimetope (YVRPLWVRME) (Pi Proteomics), 1µg/ml of 8.3 peptide (KYNKANVEL) (Chi Scientific), or 100ng/ml of OT-I peptide (SIINFEKL) (Pi Proteomics). Spleen and lymph node cells from 8 week female NOD mice were stimulated with soluble 2ug/mL anti-CD28 antibody (BioXCell) and plate-bound 2ug/mL anti-CD3 antibody (BioXCell). Beginning on day 2 post-activation, cells were maintained in media containing 10 IU/ml rhIL-2 (AIDS Research and Reference Reagent Program, Division of AIDS, NIAID, NIH from Dr. Maurice Gately, Hoffmann - La Roche Inc). Day 6-9 post-activation, BDC2.5 or 8.3 T cells were labeled with 1µM VPD (BD), 2 µM CFSE (Invitrogen), 20 µM CMTMR (Invitrogen), or 5µM eFluor670 (eBiosciences) for 25-30 minutes at 37°C or 0.5 µM CTFR (ThermoFisher Scientific) or 2.5 µM CTY (ThermoFisher Scientific) for 15 minutes at 37°C. 1×10^7^ T cells were transferred intravenously into recipients with the timing indicated in the schematics provided for each experiment.

### Generation and Treatment of BM Chimeras

#### CD11c-DTR mixed BM chimeras

Eight-week old WT NOD female mice were lethally γ-irradiated with 2 doses of 500 Rads. Following irradiation, 1×10^7^ bone marrow cells (90% NOD.CD11c-DTR and 10% NOD.8.3.CD2-dsRed.CD11c-DTR or 95% NOD.CD11c-DTR and 5% NOD.CD11c-DTR.CD2-dsRed) were injected intravenously via the tail. Bone marrow chimeras were allowed to reconstitute for at least 8 weeks prior to use. Following reconstitution mice were injected with 2 doses of 200ng diphtheria toxin (Sigma) or control PBS 24 hours apart. Twenty-four hours after the final diphtheria toxin treatment islets were isolated and mounted for 2-photon imaging.

#### ZBTB46-DTR BM chimeras

C57BL/6.RIP-mOva mice were γ-irradiated with 2 doses of 500 Rads. Following irradiation, 1×10^7^ C57BL/6.ZBTB-46-DTR bone marrow cells were injected intravenously via the tail. Bone marrow chimeras were allowed to reconstitute for at least 8 weeks prior to use. C57BL/6.OT-I.Ubiquitin-GFP CD8+ T cells were enriched from single-cell suspensions of lymph nodes and spleens using mouse CD8-negative selection kits (Stem Cell Technologies). Enriched CD8 T cells (5×10^6^) were injected into ZBTB-46-DTR-RIP-mOva bone marrow chimeras to induce islet infiltration. Two days prior to islet isolation, mice were treated with 2 doses of 200ng diphtheria toxin or control PBS 24 hours apart to deplete ZBTB-46 expressing cells. Day 5-7 post T cell transfer, islets were isolated for explanted imaging. Following islet isolation, T cell motility was assessed as by 2-photon imaging.

### Flow Cytometry: Tissue preparation and Antibodies

#### Tissue isolation and digestion

Islets were isolated and digested as described above. Pancreatic (draining) or Inguinal (non-draining) lymph nodes were isolated and shredded using needles. Lymph nodes were then digested with 0.4 Wunsch Units/ml Collagenase D (Roche) and 250 µg/ml DNAse I (Roche) in HBSS (Cellgro) with 10% FBS (Hyclone) at 37°C for 30 min.

#### MERTK identification panel

Islets were isolated from 4-18 week old NOD.CX3CR1-GFP+/- mice and dissociated as previously described. Islet samples were stained with fluorescent labeled antibodies (CD11c PE-Cy7 (Biolegend), MHCII PE (Biolegend), CD45 Pacific Blue (Biolegend), unlabeled goat α-MERTK (R&D systems), and donkey α-goat secondary (AF647, Jackson Immuno)).

#### T cell activation panel

Isolated islet and lymph node cells were surface stained with CD45 BUV395 (BD), CD4 BV711 (Biolegend), CD8 APC-780 (eBioscience), CD62L Pacblue (Biolegend), CD44 FITC (Biolegend) for 30 minutes on ice.

#### Granzyme B intracellular panel

Isolated islet and lymph node cells were stained with fluorescent antibodies (CD45 BUV395 (BD), CD4 BV711 (Biolegend), CD8 FITC (Biolegend), dump PerCP-cy5.5 (CD11c (eBioscience), CD11b (Biolegend), Ter119 (BD), F4/80 (Biolegend), and CD19 (BD)) on ice for 30 minutes. Following surface staining, samples were fixed with 1% PFA (Sigma) and 3% sucrose (Sigma) for 13 minutes. Following fixation samples were stored overnight in permeabilization buffer (eBioscience). After overnight permeabilization, samples were stained intracellularly with Granzyme B (Pacific Blue, Biolegend) for 30 minutes at room temperature.

#### APC identification panel

Islets were isolated from ZBTB46-DTR BM chimera mice dissociated as previously described. Islet samples were stained with fluorescent labeled antibodies (CD45 BUV395 (BD), CD90.2 PE-Cy7 (Biolegend), CD19 PE (Biolegend), F4/80 PerCP-Cy5.5 (Biolegend), MHCII FITC (Biolegend), CD11c BUV805 (BD), CD11b APC-Cy7 (Biolegend), XCR1 BV785

<u>(Biolegend),</u> unlabeled goat α-MERTK (R&D systems), and donkey α-goat secondary (AF647, Jackson Immuno).

### MERTK identification in CD11c-DTR-GFP mice

Islets were isolated from NOD.CD11c-DTR-GFP mice as previously described. Islet samples were stained with fluorescent labeled antibodies (CD45 BUV395 (BD), CD90.2 BUV737 (BD), CD19 BUV737 (BD), CD11c PE-Cy7, CD11b AlexaFluor647, XCR1 BV785, MHCII Biotin (BD) + Streptavidin Qdot605 (Invitrogen), and rabbit α-MERTK (Abcam) + donkey α-rabbit BV421 (Biolegend).

### Flow Cytometry and Analysis

All samples were run on an LSR Fortessa (BD) or Aurora (Cytek). Analysis and conventional compensation were performed in FlowJo (TreeStar). Spectral unmixing was performed using or SpectroFlo (Cytek).

### In vivo treatment with UNC2025

UNC2025 is a selective MERTK/Flt3 kinase inhibitor (Zhang et al., 2014)that was initially developed to inhibit MERTK signaling in MERTK+ leukemia cells (Zhang et al., 2014; DeRyckere et al., 2017). UNC2025 is orally bioavailable; however, it has a short half-life that requires twice daily dosing. UNC2025 (Meryx) was dosed at 30 mg/kg, which has been used to treat mice in a cancer setting (Cook et al., 2013; Cummings et al., 2015). Mice were treated twice daily for up to 4 weeks as indicated with daily monitoring of blood glucose and body weight.

### Diabetes incidence

For disease incidence experiments all mice had normal glycemia prior to treatment. Blood glucose readings were taken daily prior to afternoon treatment. Mice with two consecutive readings >250 mg/dl were considered diabetic, and were euthanized. Mice were treated for at least 2 weeks unless sacrificed due to diabetes.

### T cell activation

WT NOD 12-16 week old female mice were treated with 30 mg/kg UNC2025 or vehicle. Sixteen hours later mice were treated a second time with UNC2025 or vehicle. One hour after final treatment islets, draining pancreatic lymph node, and non-draining inguinal lymph node were harvested and digested as described above. Single cell suspensions were labeled with the T cell activation panel described above.

### In vivo granzyme B

WT NOD 12-16 week old female mice were treated for 16 hours with UNC2025 or vehicle in the same manner as described above. At the same time as the final UNC2025 treatment, mice were injected intravenously with 250 µg of Brefeldin A (Sigma). Four hours after Brefeldin A injection, islets, draining pancreatic lymph node, and non-draining inguinal lymph node were harvested. Samples were maintained in 10 µg/ml Brefeldin A on ice when possible until fixation. Islets and lymph nodes were digested and dissociated as described above.

Staining was performed using the Granzyme B panel as described above. Due to changes in cytometer settings between experiments, mean fluorescent intensity (MFI) data were normalized for the purposes of graphing all experiments together. All samples were normalized to the average value of the vehicle controls from the same tissue in the same experiment. However, statistical analysis was performed on the data prior to normalization.

### Islet section scoring

WT NOD female mice greater than 20 weeks of age were treated twice daily with 30mg/kg UNC2025 or vehicle. For islet sections, mice were treated twice 16 hours apart with UNC2025 or vehicle. One hour after the final treatment, the whole pancreas was excised and fixed in formalin (VWR) for 4 days. After fixation samples were stored in 70% ethanol until embedding in paraffin. Seven-µm sections were cut and fixed with Hematoxylin and Eosin. Embedding, sectioning and staining were performed by the University of Colorado Denver Anschutz histology core. Section scoring was performed blindly on a scale of 0 (no infiltration) to 4 (completely infiltrated).

Formalin-fixed paraffin embedded human pancreas sections were obtained from nPOD. Sections were deparaffinized using mixed xylenes (Sigma-Aldrich), and antigen retrieval was done using Tris pH10 antigen retrieval buffer in a pressure cooker for 10 minutes. Islets were stained for insulin (polyclonal guinea pig), CD3 (clone CD3-12), MERTK (clone Y323), and CD68 (clone KP1), all from Abcam. Secondaries included anti-GP-Cy3, anti-rat-DL649, anti-Mouse-biotin, anti-rabbit-biotin (all Donkey antibodies from Jackson Immuno) and Streptavidin-FITC, Streptavidin-BV421 (Biolegend). MERTK & CD68 stains were done sequentially with biotin-avidin blocking (Life Technologies) between stains. Differential staining patterns for CD68 and MERTK confirmed successful blocking. Approximately 10 insulin-containing islets were imaged per donor using a Zeiss 710 confocal microscope. Islet images were analyzed in a blinded manner using Imaris software.

### Analysis of UNC2025 treatment by imaging

Activated BDC-2.5 and 8.3 T cells were labeled with VPD or eFluor670 and transferred IV into 12-16 week old NOD.CD11c-mCherry female mice. Eight hours following T cell transfer mice were treated with 30 mg/kg UNC2025 or vehicle. Sixteen hours later mice were again treated with UNC2025 or vehicle. One hour later islets were isolated and imaged using our explanted imaging technique as previously described (explanted islet imaging).

Activated OT-I.Ub-GFP T cells were transferred into WT or MERTK-/- tumor bearing hosts 5-7 days before treatment. 18h prior to tumor harvest, mice were treated with 30 mg/kg UNC2025 or vehicle by oral gavage. Whole tumors were excised and mounted on cover slips for imaging. To ensure comparable microenvironments, tumor imaging was done in superficial regions of the tumor, where OT-I T cells were present, and the collagen rich tumor capsule was visible.

UNC2025 treated tissue explants were maintained in 50 μM UNC2025 for the duration of isolation and imaging. Imaging duration was 30 minutes long; image analysis was performed using Imaris and Matlab as previously described.

### Analysis of T cell motility within MERTK-/- islets

For studies involving NOD.MERTK-/- mice, which do not spontaneously develop diabetes, islet infiltration was induced in NOD.MERTK-/- or control WT mice by transferring 1×10^7^ activated BDC-2.5 T cells 3-6 days prior to dye-labeled T cell transfer. Following initiation of islet infiltration 1×10^7^ VPD or eFlour670 dye labeled activated BDC2.5 and 8.3 T cells were transferred intravenously. Twenty-four hours following labeled T cell transfer islets were isolated and imaged by 2-photon microscopy. Imaging duration was 30 minutes long; image analysis was performed using Imaris and Matlab as previously described.

### Immunofluorescent staining and analysis of human islets

Deidentified formalin-fixed paraffin-embedded pancreas sections were obtained from the nPOD consortium. The experiments were determined to be non-human subject research by the National Jewish Health Institutional Review Board. Donor information is noted in Table S1.

Sections were deparaffinized with xylenes and antigen retrieval was performed with Tris pH10 buffer in a pressure cooker. Sections were blocked with 10% donkey serum (Jackson ImmunoResearch) and a biotin-avidin blocking kit (Thermo Fisher). Pancreas sections were then stained for Insulin, MERTK, CD68, and CD3 (Abcam: ab955, ab52968, ab7842, ab11089). Anti-insulin and anti-CD3 were detected with fluorescently labeled secondary antibodies (Jackson Immunoresearch: 706-166-148, 712-606-153). CD68 was detected using a biotin-conjugated secondary antibody (Jackson Immunoresearch: 715-065-151) followed by fluorescently conjugated streptavidin (Biolegend 405201). The avidin-biotin blocking kit was then used to block any free streptavidin and biotin. After blocking of CD68-biotin-avidin was complete, MERTK was then detected using a biotin-conjugated secondary antibody (Jackson Immunoresearch: 711-065-152) followed by fluorescently conjugated streptavidin (Biolegend 405226). Data were acquired using a Zeiss LSM700 microscope. 10 pancreas images from each donor were acquired based on expression of insulin.

Samples were blinded prior to analysis and analysis was performed by a third party. 10-15 islets per donor were analyzed, with an average of 13.10 islets per Control donor and 13.11 islets per T1D donor analyzed. Image visualization and determination of islet area were performed using Imaris (Oxford Instruments).

### Online supplemental material

Table S1 displays information about the donor samples analyzed in Figure 8. Figure S1 shows that the CD11c-DTR-GFP cells are effectively depleted in the islets upon DT treatment, and that the islet CD11c-DTR-GFP cells include CD11b+, XCR1+ and MERTK+ populations. Figure S2 shows that Zbtb46-expressing dendritic cells are not the subset of CD11c+ cells in the islets that prevent T cell arrest. Figure S3 shows that MERTK deficiency or MERTK inhibition do alter the mononuclear phagocyte frequency in the islets. Figure S4 shows that costimulatory and coinhibitory molecule expression on CD11c+ cells is not altered by MERTK inhibition. Figure S5 shows examples of CD44/CD62L flow cytometry gating. Video S1 is an example of the data analyzed in Figure 1, which shows that T cell motility in the islets is reduced upon CD11c+ cell depletion. Video S2 is an example of the data analyzed in Figure 3, which shows that T cell motility is reduced in MERTK deficient islets. Video S3 is an example of the data analyzed in Figure 5, which shows that T cell motility on the surface of a melanoma tumor is reduced with either MERTK-inhibition or MERTK-deficiency in the tumor host. Video S4 is an example of the data analyzed in Figure 4 and Figure 6, which shows that in the islets, sustained interactions between islet antigen specific T cells and CD11c+ cells are increased with MERTK inhibition, while T cells that are not specific for islet antigen show now change in T cell-CD11c+ cell interactions.

## Author Contributions

Conceptualization, R.S.F and R.S.L.; Methodology, R.S.F., R.S.L., J.C.W., and J.J.; Investigation, R.S.L., J.C.W., K.E.D., E.R., A.M.S., D.T., S.F.Y., B.N.B., and R.S.F.; Analysis, R.S.L., R.S.F., J.C.W., E.R., B.N.B, and J.J.; Writing – Original Draft, R.S.F. and R.S.L.; Writing–Review & Editing, R.S.F., R.S.L., J.C.W., A.M.S., and J.J.; Funding Acquisition, R.S.F.; Supervision, R.S.F.

## Acknowledgements

We thank Meryx Inc. for providing UNC2025; Dr. Jonathan Katz, Dr. Katie Haskins, Dr. Qizhi Tang, and Dr. Roland Tisch for mice; Dr. Katie Haskins, Dr. John Cambier, Dr. Michael Holers, Dr. Ron Gill, Dr. Claudia Jakubzick, Dr. Ross Kedl, Dr. Rocky Baker, Dr. Mira Estin, and Dr. Scott Thompson for guidance and helpful scientific input; Dr. Pippa Marrack, Dr. John Kappler, Dr. Ross Kedl, Dr. Claudia Jakubzick, Dr. Larry Wysocki, and Dr. Laurel Lenz for reagents; Robert Long, Matthew Gebert, Brianna Traxinger, Marlie Fisher, Katie Morgan, and Jessica Olivas, Scott Beard, the National Jewish Health Flow Cytometry Core Facility, the National Jewish Health Biological Resource Center, the Barbara Davis Center Flow Cytometry Core, and University of Colorado Vivarium for animal husbandry and technical assistance; Bonnie Levitt and Joseph Kleponis for assistance with coding MATLAB analysis scripts. This research was performed with the support of the Network for Pancreatic Organ donors with Diabetes (nPOD; RRID:SCR_014641), a collaborative type 1 diabetes research project sponsored by JDRF (nPOD: 5-SRA-2018-557-Q-R) and The Leona M. & Harry B. Helmsley Charitable Trust (Grant#2018PG-T1D053). Organ Procurement Organizations partnering with nPOD to provide research resources are listed at http://www.jdrfnpod.org//for-partners/npod-partners/. This work was supported by funding from JDRF #2-2012-197 (RSF), the William & Ella Owens Medical Research Foundation (RSF), NIH 1R01DK111733-01 (RSF), George S. Eisenbarth nPOD Award from JDRF/Helmsley Trust (RSF), CRI #AWD-112499 (RSL), T32: 5T32AI007405-27 (JCW) and the University of Colorado Diabetes Research Center NIH P30-DK116073.

## Abbreviations

T1D: Type 1 diabetes

APCs: Antigen presenting cells

DCs: Dendritic cells

BM: Bone marrow

DT: Diptheria toxin

mOva: membrane bound ovalbumin

RIP: Rat insulin promotor

nPOD: network of pancreatic organ donors with diabetes

MFI: Mean fluorescent intensity

PD-1: Programmed death 1

PD-L1: Programmed death-ligand 1

TAM: TYRO3 AXL MERTK family members

PBS: Phosphate buffered saline

HBSS: Hanks’ balanced salt solution

FBS: Fetal bovine serum

NOD: Non-obese diabetic

TCR: T cell receptor

Tregs: Regulatory T cells

PLN: Pancreatic Lymph Node

ILN: Inguinal Lymph Node

---

**Table S1:** nPOD donor information

| Disease | ID | Sex | Age | Ethnicity | Cause of death | T1D Duration (years) | Auto-antibody | C-peptide | BMI |
| --- | --- | --- | --- | --- | --- | --- | --- | --- | --- |
| Control | 6339 | Male | 23.3 | Caucasian | Head trauma | 0 | Negative | 10.56 | 25 |
| Control | 6178 | Female | 24.5 | Caucasian | Anoxia | 0 | Negative | 4.55 | 27.5 |
| Control | 6331 | Female | 27 | African American | Anoxia | 0 | Negative | 3.01 | 24 |
| Control | 6333 | Female | 27.1 | Caucasian | Anoxia | 0 | Negative | 9.37 | 24.9 |
| Control | 6254 | Male | 38 | Caucasian | Anoxia | 0 | Negative | 6.43 | 30.5 |
| Control | 6425 | Female | 38.66 | Caucasian | Anoxia | 0 | Negative | 8.06 | 28.28 |
| Control | 6015 | Female | 39 | Caucasian | Anoxia | 0 | Negative | 1.99 | 32.2 |
| Control | 6250 | Male | 40 | Caucasian | Head trauma | 0 | Negative | 7.31 | 27.9 |
| Control | 6097 | Female | 43.1 | Caucasian | Stroke | 0 | Negative | 16.76 | 36.4 |
| Control | 6369 | Male | 44 | Caucasian | Stroke | 0 | Negative | 6.42 | 18.8 |
| T1D | 6051 | Male | 20.3 | Caucasian | Head trauma | 13 | mIAA | <0.05 | 21.5 |
| T1D | 6435 | Female | 24.75 | African American | Anoxia | 14.75 | Negative | 0.09 | 26.9 |
| T1D | 6196 | Female | 26.5 | African American | Anoxia | 15 | GADA, mIAA | 0.48 | 26.6 |
| T1D | 6039 | Female | 28.7 | Caucasian | Head trauma | 12 | GADA, IA-2A, ZnT8A, mIAA | <0.05 | 23.4 |
| T1D | 6038 | Female | 37.2 | Caucasian | Anoxia | 20 | Negative | 0.2 | 30.9 |
| T1D | 6328 | Male | 39 | Hispanic | Anoxia | 20 | GADA, mIAA | <0.02 | 24 |
| T1D | 6438 | Female | 39.01 | Caucasian | Anoxia | 10 | GADA | <0.02 | 35 |
| T1D | 6175 | Male | 42 | Caucasian | Stroke | 12 | GADA | 3.48 | 19.8 |
| T1D | 6307 | Female | 45 | Caucasian | Anoxia | 10 | GADA, mIAA | <0.05 | 19.5 |

## Supplemental Figure Legends

Figure S1: CD11c-DTR-GFP cells are effectively depleted in the islets and include CD11b+, XCR1+ and MERTK+ populations. NOD.CD11c-DTR-GFP bone marrow (BM) chimeras containing NOD.CD11c-DTR-GFP BM were treated twice with PBS or Diphtheria Toxin (DT). Islets from **(A)** treated NOD.CD11c-DTR BM chimeras or **(B)** NOD.CD11c-DTR-GFP+CD2.dsRed+ female mice were isolated, digested and dissociated, stained, and analyzed by flow cytometry. **(A)** Quantification of CD11c+ cells in islets following PBS or DT treatment. **(B)** Quantification of the composition of the CD11c-DTR-GFP+ versus GFP-mononuclear phagocytes in the islets. **(A)** n = 12 mice (PBS), n = 16 mice (DT) from 7 independent experiments; Error bars = SEM. Statistics: Students T-test, ***p<0.001. **(B)** n=4 mice from 3 independent experiments.

Figure S2: Zbtb46-expressing dendritic cells are not the subset of CD11c+ cells in the islets that prevent T cell arrest. (A-D) Islet infiltration was induced in B6.Zbtb46-DTR.RIP-mOva BM chimeras by the transfer of naïve OT-I.GFP T cells. **(A)** Three to five days after T cell transfer, the chimeras were treated twice with PBS (Control) or DT (Zbtb46-depleted). 48h post-treatment initiation, T cell motility was analyzed in explanted islets by 2-photon microscopy. **(B)** Average T cell crawling speed per islet. **(C)** Arrest coefficient: % of time a T cell is moving <1.5 µm/min. **(D)** Average islet infiltration levels. **(E)** Flow cytometric analysis of mononuclear phagocytes in the islets of PBS (Control) or DT (Zbtb46-depleted) treated NOD.Zbtb46-DTR BM chimeras, 48h post-treatment initiation. **(A-D)** Data pooled from 3 independent experiments. n = 10 islets from PBS treated mice and 26 islets from DT treated mice. **(E)** Data pooled from 4-5 mice per condition in 2 independent experiments. **(B,D,E)** Statistics: Students T test. **(C)** Statistics: 2-way ANOVA. Error bars: SEM. **(B-E)** Error bars: SEM, *p<0.05.

Figure S3: Mononuclear phagocyte frequency in the islets is not altered by MERTK deficiency or MERTK inhibition. (A) Cell frequency of WT and MERTK-/- NOD mice was quantified in the islets by flow cytometry. Statistics: Two-tailed Student t test. n = 3 experiments. Error bars: SEM. **(B)** WT NOD female mice were treated with 30mg/kg UNC2025 or saline vehicle twice daily by oral gavage for 4 weeks. Cell frequency was quantified in the islets by flow cytometry. Statistics: Two-tailed Student t test. n = 6-8 mice per group from 2 experiments. Error bars: SEM.

Figure S4: MERTK inhibition does not alter the expression of islet costimulatory or coinhibitory ligands. WT NOD female mice were treated with 30mg/kg UNC2025 or saline vehicle for 18h. Islets, pancreatic lymph nodes, and inguinal lymph nodes were then harvested, dissociated, antibody stained, and analyzed by flow cytometry. Data show expression on CD45+CD11c+ cells. n=3-8 mice from 2-4 experiments. FMO = fluorescence minus one, unstained gMFI of gated population.

Figure S5: Examples of CD44/CD62L flow cytometry plots after vehicle or UNC2025 treatment. Data quantified in Figure 7A. WT NOD female mice were treated with 30mg/kg UNC2025 or vehicle twice by oral gavage. Islets and lymph nodes were isolated 17h post-treatment initiation, digested, stained and analyzed by flow cytometry. Gating of Islet samples was done independently from gating of lymph node samples due to partial cleavage of CD44 by Collagenase P, which was used for the islet isolation. Previous tests treating split samples of splenocytes with Collagenase P under the conditions used for islet isolation compared to no Collagenase P showed diminished florescence intensity of CD44, but equal frequencies when gating was adjusted. CD44 gates were set for islet samples and lymph node samples using CD62L+ cells as a CD44-population.

## Movie Legends

Movie 1: **CD11c+ cells in the islets prevent T cell arrest.** NOD.CD11c-DTR bone marrow (BM) chimeras containing 90% NOD.CD11c-DTR BM +10% NOD.CD11c-DTR.8.3.CD2-dsRed BM were treated twice with PBS (Intact) or DT (CD11c depleted). 48h post-treatment initiation, T cell motility was analyzed in explanted islets by 2-photon microscopy. Representative islet movies from PBS control or DT treated mice with 10 minute tracks of motion. 8.3 T cells are shown in white and CD11c-DTR-GFP is shown in green. Dashed line depicts islet border. Time (min:sec); scale bar= 50µm. Representative of data quantified in Figure 1A-G.

Movie 2: **Islet antigen-specific T cells in MERTK-/- islets show decreased motility and increased T cell arrest.** Activated BDC-2.5 CD4 T cells were transferred into NOD or NOD.MERTK-/- female mice to initiate islet infiltration. Fluorescently labeled activated 8.3 CD8 T cells (white) and BDC-2.5 CD4 T cells (red) were then transferred into the same mice 3-6 days later. Twenty-four hours after fluorescent T cell transfer, islets were isolated and analyzed by 2-photon microscopy. Representative islet movies from WT NOD or NOD.MERTK-/- mice with 10 minute tracks of motion. Dashed line depicts islet border. Time (min:sec); scale bar= 50µm. Representative of data quantified in Figure 3.

Movie 3: **MERTK signaling prevents T cell arrest in a solid tumor model.** B78ChOva tumor cells were implanted in C57BL/6 or C57BL/6.MERTK-/- mice. Activated OT-I.Ub-GFP CD8 T cells (green) were then transferred into tumor bearing mice mice 4-6 days later. Five to seven days after fluorescent T cell transfer, mice were treated with UNC2025 or vehicle. 18h after treatment, tumors were excised and analyzed by 2-photon microscopy. Representative movies of OT-I T cell (green) motility on the tumor surface with 10 minute tracks of motion. Collagen (blue) delineates the tumor surface. Representative of data quantified in Figure 5.

Movie 4: **MERTK signaling prevents islet T cell-APC interactions in an antigen-dependent manner.** Fluorescently labeled, activated 8.3 CD8 T cells (white), BDC-2.5 CD4 T cells (red), and polyclonal T cells (blue) were co-transferred into female NOD.CD11c-YFP (green) mice. Mice were treated twice daily with saline vehicle or 30 mg/kg UNC2025, a small molecule inhibitor of MERTK and Flt3. 16h following the initial treatment, islets were isolated and analyzed by 2-photon microscopy. Representative islet movies from vehicle or UNC2025 treated mice with 5-minute tracks of motion and circles highlighting sustained interactions between T cells and APCs. Time (min:sec); scale bar= 30µm. Representative of data quantified in Figure 4 and Figure 6.

