## Supplemental Figure S1 for "MERTK on mononuclear phagocytes regulates T cell antigen recognition at autoimmune and tumor sites"

### Figure S1: CD11c-DTR-GFP cells are effectively depleted in the islets and include CD11b+, XCR1+ and MERTK+ populations

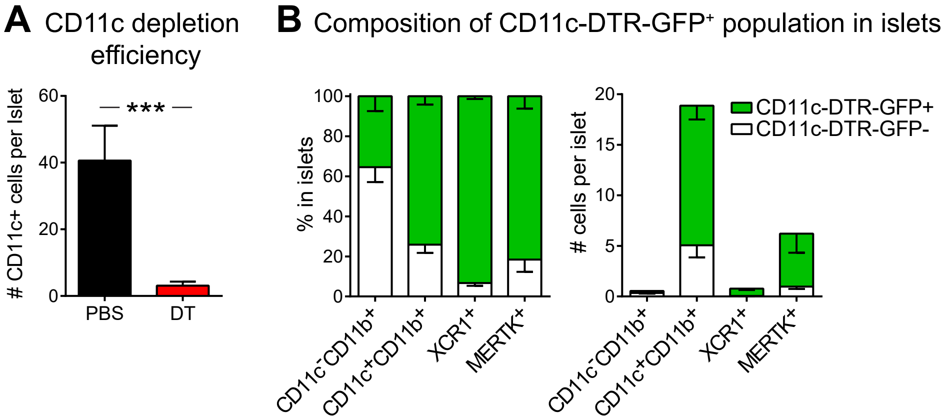
