## Supplemental Figure S2 for "MERTK on mononuclear phagocytes regulates T cell antigen recognition at autoimmune and tumor sites"

### Figure S2: Zbtb46-expressing dendritic cells are not the subset of CD11c+ cells in the islets that prevent T cell arrest.

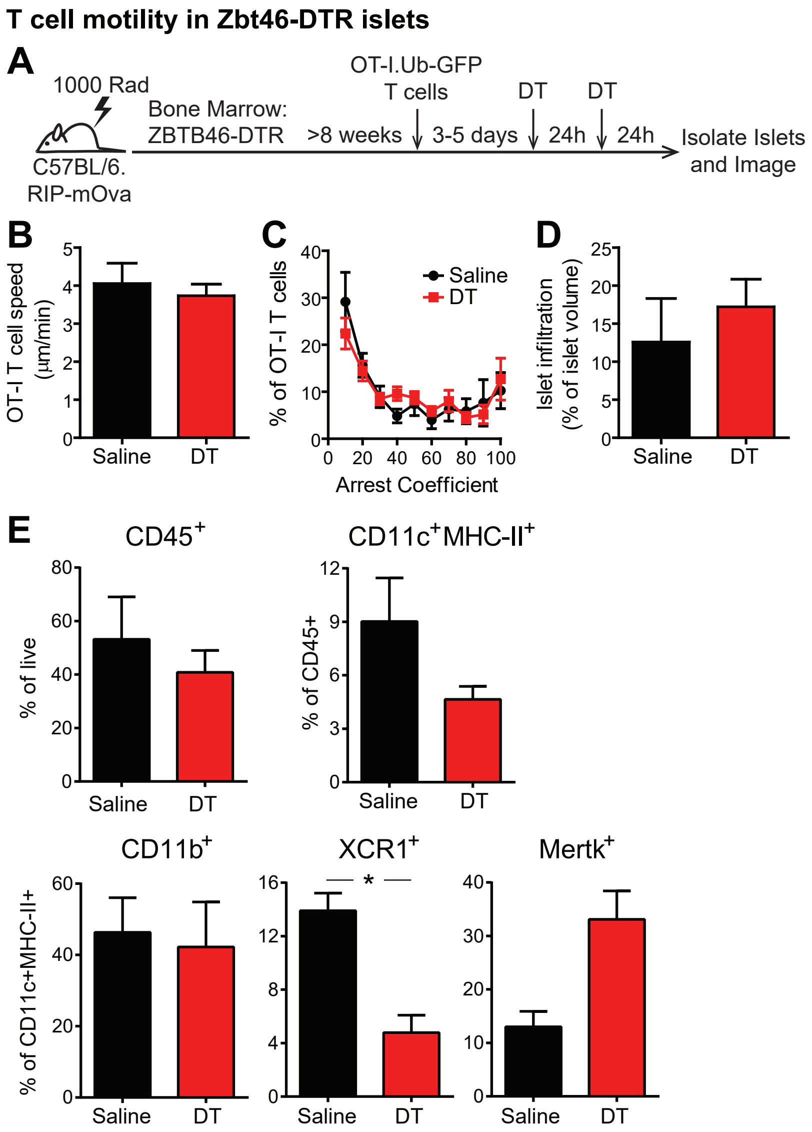
