## Supplemental Figure S3 for "MERTK on mononuclear phagocytes regulates T cell antigen recognition at autoimmune and tumor sites"

### Figure S3: Mononuclear phagocyte frequency in the islets is not altered by MERTK deficiency or MERTK inhibition

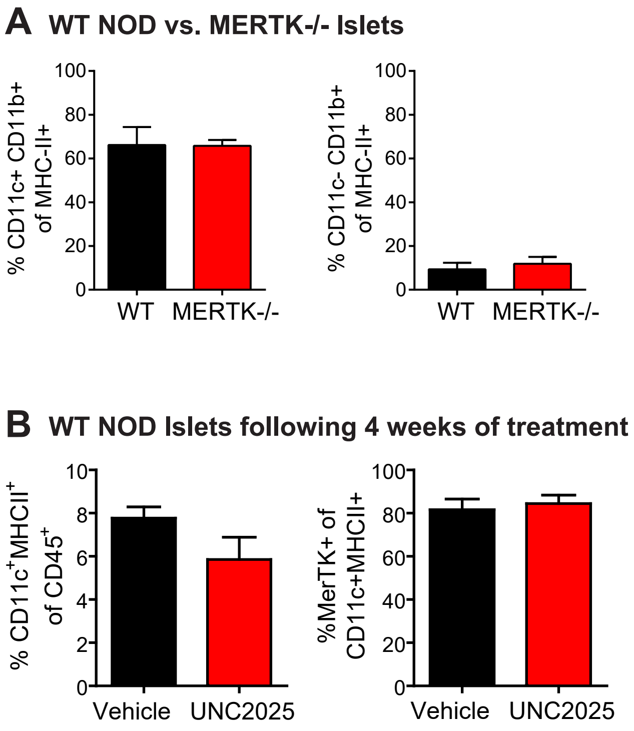
