## Supplemental Figure S4 for "MERTK on mononuclear phagocytes regulates T cell antigen recognition at autoimmune and tumor sites"

### Figure S4: MERTK inhibition does not alter costimulatory molecule expression on islet CD11c+ cells

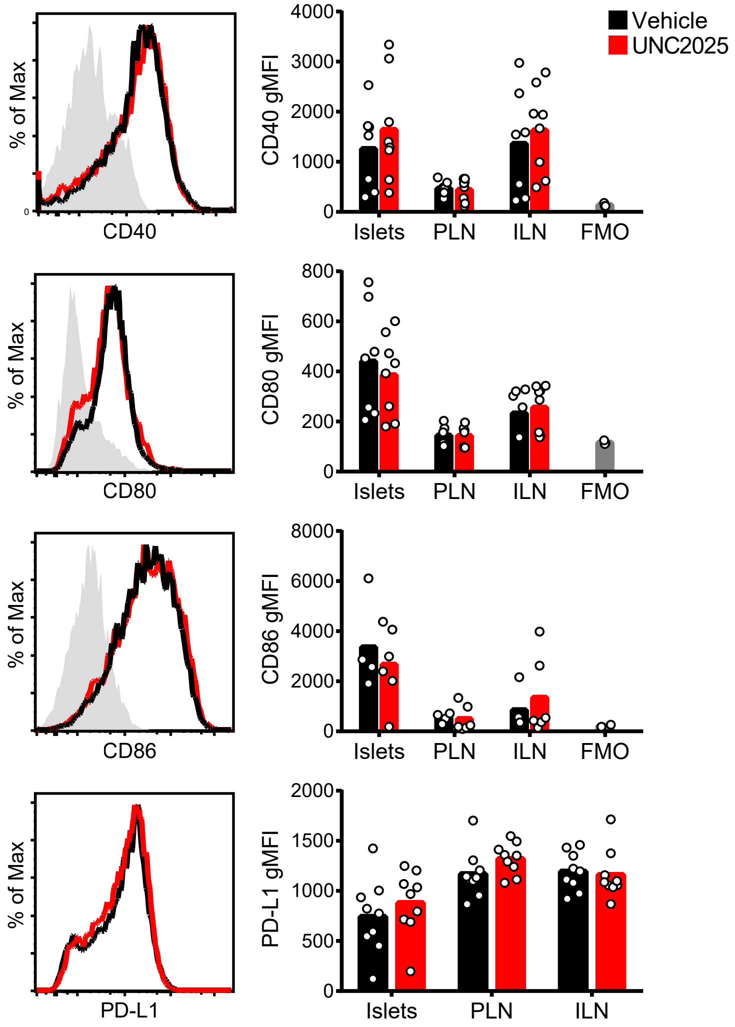
