## Supplementary figures and images for "MERTK on mononuclear phagocytes regulates T cell antigen recognition at autoimmune and tumor sites"

### Supplemental Figure S5

#### Figure S5: Examples of CD44/CD62L flow cytometry plots after vehicle or UNC2025 treatment

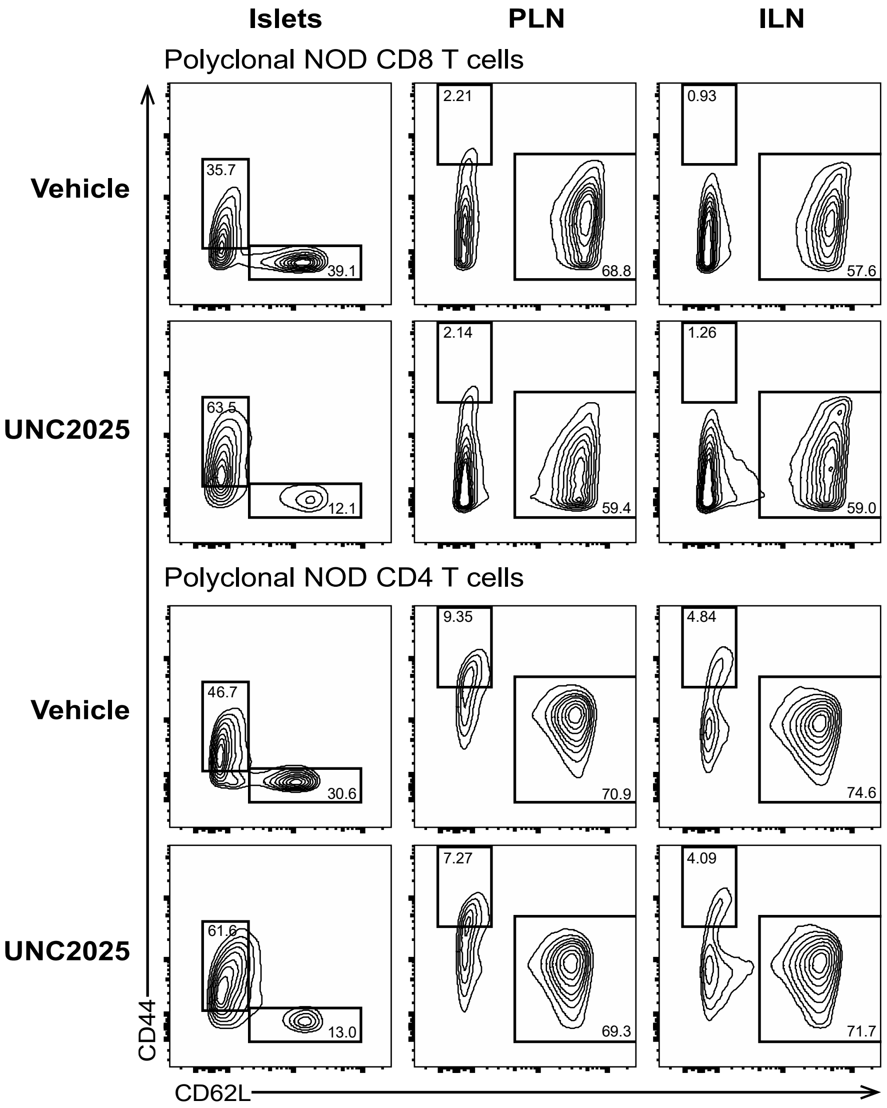
